# eCV: Enhanced coefficient of variation and IDR extensions for reproducibility assessment of high-throughput experiments with multiple replicates

**DOI:** 10.1101/2023.12.18.572208

**Authors:** Agustin Gonzalez-Reymundez, Kylie Shen, Wayne Doyle, Sichong Peng, Kasey Hutt, Stephanie Bruns

## Abstract

**Motivation:** Reproducibility assessment is essential in extracting reliable scientific insights from highthroughput experiments. Inconsistency between technical replicates poses a challenge, particularly clear in next generation sequencing technologies based on immunoprecipitations, where the assessment of reproducibility in peak identification is a critical analytical step. While the Irreproducibility Discovery Rate (IDR) method has been instrumental in assessing reproducibility, its standard implementation is constrained to handling only two replicates. In the current era of steadily growing sample sizes, eased by multiplexing and reduced sequencing costs, highly performing methods that handle any number of replicates are desirable.

**Results:** We introduce three novel methods for reproducibility assessment in high-throughput data that handle an arbitrary number of replicates. The first, general IDR (gIDR), extends the standard IDR by adapting its Expectation-Maximization (EM) algorithm to handle distributions of any dimensions dictated by the number of replicates. The second, meta-IDR (mIDR), employs a meta-analysis approach, calculating local IDR scores for all pairs of replicates and combining them using standard probability rules. The third method introduces an “enhanced” Coefficient of Variation (eCV), ranking features based on intensity and variability, using a parametric bootstrap approach to obtain an index analogous to local IDR. Comparative analysis with traditional IDR in simulated and experimental data reveals the heightened performance of the proposed multivariate alternatives under varying scenarios, thereby addressing the critical challenge of reproducibility assessment in contemporary high-throughput experiments.

**Availability and implementation:** The described methods are implemented as an R package: https://github.com/eclipsebio/eCV

## 1 INTRODUCTION

The reproducibility of high-throughput experiments is essential in assuring high-quality results and valid scientific insights (Nicholson and Holmes, 2017). Lack of consistency within technical replicates, whether coming from heightened variability or diminished mean intensities, can undermine the reliability of the finding. For example, the lack of reproducibility between peaks in separated ChIP-seq libraries has been recognized as one of the most important pitfalls of the technology (Mundade *et al*., 2014; Landt *et al*., 2012). Li et al. (2011) introduced the Irreproducibility Discovery Rate (IDR) to measure the reproducibility of high-throughput data. This method robustly models the distribution of omic feature intensity along each technical replicate using a copula model (CM), and models conveniently the joint distribution of omic features in each sample replicate based only on the marginal distributions (Genest and Nešlehová, 2013).

Within the IDR framework, features are transformed and assumed to come from a mixture of two normal distributions, each representing one of two possible states: reproducible or irreproducible. An expectation-maximization (EM) algorithm (Dempster et al., 1977) is then applied to estimate the parameters of this mixture, including the mean and variance of each normal distribution along with the mixing probability. Upon convergence, the likelihood of an omic feature belonging to the reproducible state is used to compute a “local” IDR score. To account for situations where the distribution of omic-feature values is unknown or too complex to approximate via elemental distributions (e.g., normal, negative binomial, gamma), a “semiparametric” CM approach is used, where marginal empirical cumulative distributions of each replicate are fed into the CM (Tsukahara, 2005). The actual IDR is calculated analogously to an adaptation step-up procedure to control the false discovery rate (FDR) (Sun and Cai, 2007; Li *et al*., 2011). After the model is fitted, features with small IDR scores are taken as “reproducible,” while features with higher IDR scores are excluded as “irreproducible.”

IDR has shown robust performance in extensive simulations (Li et al., 2011) and has also been adopted by the ENCODE project as the gold standard to assess the reproducibility of ChIP-seq data (Landt *et al*., 2012). Nevertheless, the classical implementation of IDR has the limitation of being unable to handle more than two replicates. The reason for this constraint imposed on IDR is the conclusion from early ChiP-seq experiments that more than two replicates would increase the assay’s cost while not significantly improving performance values, such as site discovery (Rozowsky *et al*., 2009). However, multiplexing and the continuous drop in sequencing costs make it more affordable for researchers to increase sample size (Chen *et al*., 2012). Larger sample sizes can improve estimates of the variability associated with noise and artifacts effects and increase statistical power (Li *et al*., 2017).

Several alternatives to measure reproducibility in an arbitrary number of replicates have been proposed, including pooling of all replicates (Chen *et al*., 2011; Landt *et al*., 2012), selection of the best pairs of replicates (Revilla-i-Domingo *et al*., 2012), or “majority rule” (Haecker *et al*., 2012). However, as Yang et al. (2014) discussed, the quality of these data-combination strategies may be susceptible to outliers, diminish signal-to-noise ratios, or compromise subsequent quantitative comparisons among samples.

To overcome these limitations, reproducibility assessment methods exploiting the true multivariate nature of multi-replicate experiments are needed. Here, we present three novel approaches to generalize the reproducibility assessment of high throughput data on an arbitrary number of replicates. The first one, general IDR (gIDR), expands the classic IDR by extending its EM approach to handle distributions of arbitrary dimensions dictated by the number of replicates. The second one called meta-IDR (mIDR), resembles a meta-analysis procedure where local IDR scores are calculated for every pair of replicates and combined using standard probability rules. The third method is based on the coefficient of variation rather than IDR. It ranks features in terms of their intensity and variability and uses a parametric bootstrap approach to estimate an index analogous to the local IDR. This method is an “enhanced” Coefficient of Variation (eCV).

All three methods presented here were compared against traditional IDR in simulated and experimental data of varying numbers of replicates. Simulated data followed three scenarios used in the original IDR publication, demonstrating situations where the reproducible features are a) many and strongly correlated, b) a few and weakly correlated, and c) even fewer but strongly correlated. The methods were also compared in miR-eCLIP (Manakov *et al*., 2022) and RBP-eCLIP (Van Nostrand *et al*., 2016) data. We show that IDR is outperformed by multivariate alternatives, particularly by gIDR when many highly correlated genuine features are present (such as miR-eCLIP, where RNAs form chimeras with highly abundant miRNAs). Methods eCV and mIDR, on the other hand, performed well when a small number of highly correlated reproducible features were present (e.g., reads clustering at transcription factor or RNA binding protein binding sites). In addition, the eCV method resulted in the most stable performance across all scenarios when a liberal threshold on the probability is used (50%). All approaches are implemented in the R package *eCV*.

## 2 MATERIALS AND METHODS

### 2.1 Data sets

#### 2.1.1 Simulated data sets

Simulated data was generated using an extension of the copula mixture model presented in Li et al. (2011), assuming 1,000 omic features with values coming from two, three, and four replicates, respectively. In each simulation, the values of the *i*-th feature were defined as *y*_*ij*_ = −log 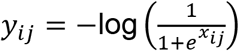. The vector holding the values of feature *i*, ***x***_*i*_ = [*x*_*i*1_, …, *x*_*i*r_], is assumed as coming from a mixture of two multivariate normal distributions (*ϕ*) of dimension r:

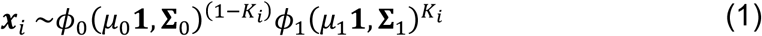

Such as

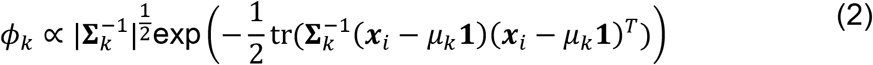

Where the dummy variable *K*_*i*_, standing for the reproducible or irreproducible classes, was sampled from a Bernoulli variable *K*_*i*_ with probability π.

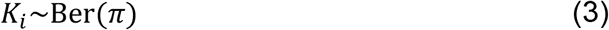

Parameters µ_*k*_ and **∑**_K_ represent a mean scalar and an rxr variance-covariance matrix, respectively. The quantity 1 was a vector of ones of length r. When *K*_*i*_ = 0, irreproducible features values were sampled from ϕ_0_, with µ_0_ = 0 and ∑_0_ = **I** being the identity matrix (i.e., a matrix with all zeros, except for ones on the diagonal). Whenever *K*_*i*_ = 1, features values were sampled from *ϕ*_1_.

In all cases, features were assumed to have unit variance, the same mean intensity (µ_1_= 2.5) and the same correlation values. This implied following a compound-symmetric model for the variance-covariance matrix 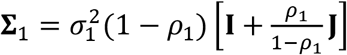, where 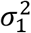and *ρ*_1_ are the reproducible class variance and correlation, respectively, **I** is an rxr identity matrix and **J** is an rxr matrix with all ones.

A summary of each parameter’s value defining each simulation scenario is presented in Table 1. Scenario 1 corresponded to a case where most features came from the reproducible class (*π* = 0.65) and were highly correlated (*ρ*_1_ = 0.84). Scenario 2 corresponded to a case with few (*π* = 0.25) lowly correlated (*ρ*_1_ = 0.40) reproducible features. Scenario 3 represented cases with even fewer (*π* = 0.05) but highly correlated (*ρ*_1_ = 0.84) reproducible features. The instructions to generate the simulated data can be found in Supplementary File S1.

**Table 1:** Parameters values defining each simulation scenario. Each simulated scenario (**Scenario 1-3**) was defined by assigning values to population parameters (**Parameter values**); proportion (***π***), mean intensity (***μ***^***1***^), variance (***σ***^***21***^), and correlation (***ρ***^***1***^) of reproducible features.

| Parameters values | Scenario 1 | Scenario 2 | Scenario 3 |
| --- | --- | --- | --- |
| $\pi$ | 0.65 | 0.25 | 0.05 |
| $\mu_1$ | 2.50 | 2.50 | 2.50 |
| $\sigma_1^2$ | 1.00 | 1.00 | 1.00 |
| $\rho_1$ | 0.84 | 0.40 | 0.84 |

#### 2.1.2 Experimental data sets

Experimental data came from two miR-eCLIP (Manakov *et al*., 2022) and one RBFOX2 eCLIP (Van Nostrand *et al*., 2016) data sets. The first scenario corresponded to miR-eCLIP on HEK293T cells transfected with two microRNAs (miRNA), *hsa-miR-1-3p* (miR-1) and *hsa*-*miR-124-3p* (miR-124). The data was arranged as intensity of messenger RNA and miRNA interactions (mRNA:miRNA), in the form of chimeric reads along two technical replicates. In this scenario, chimeric peaks annotated with the transfected miRNA (i.e., mRNAs forming chimeras with either one of the exogenous miRNAs) were used to measure performance. In this context, peaks annotated with the transfected miRNA and classified as reproducible, were taken as “true positives”. Meanwhile, endogenous mRNA:miRNA chimeras, classified as irreproducible, were taken as “true negatives”. We justified this choice by the fact that most endogenous chimeras in this data set had significantly lower intensities than the ones involving transfected miRNAs (Supplementary File S2). Since only two replicates were available (r = 2), the only relevant comparisons were between IDR and eCV, since both gIDR and mIDR converge exactly to IDR when r = 2.

The second experimental scenario also consisted of miR-eCLIP data, but this time on wild-type K562 cells, using two, three, and four replicates. Since all chimeras involved endogenous RNAs, we used a different definition of “truth”. In this scenario, true positives were defined by chimeras with positive seed match on the 3’ untranslated region (UTR). A positive seed match (i.e., high-quality alignment between the seed region of a miRNA and the complementary sequence of its target mRNAs) on the 3’UTR of a gene (the region of the targeted gene most likely to generate stable and effective RNA duplexes) show a potential regulatory interaction (Gu *et al*., 2009). Seed matching analysis was performed with *scanMiR* (v4.2.1) and *scanMiRData* (v4.2.1) (Soutschek *et al*., 2022).

The third experimental scenario comprised RBFOX2-eCLIP data, organized as peaks generated around clusters created by crosslinking and immunoprecipitating RBFOX2 across two, three, and four technical replicates from wild-type HEK293T cells. Peaks enriched with the RBFOX2 motif were detected using *HOMER* (Heinz *et al*., 2010). Peaks enriched with the RBFOX2 motif and mapping onto introns or the 3’UTR of targeted transcripts were taken as proxies of “truly reproducible” features. Every other type of peak was considered “truly irreproducible.” A summary of the characteristics of each experimental scenario is provided in Table 2.

**Table 2:** Characteristics of each experimental data scenario.

| Characteristics | Scenario 1 | Scenario 2 | Scenario 3 |
| --- | --- | --- | --- |
| **Technology** | miR-eCLIP | miR-eCLIP | RBP-eCLIP |
| **Target of immune precipitation** | AGO2 | AGO2 | RBFOX2 |
| **Experimental condition** | Transfections with miR-1 and miR-124 | Wild type | Wild type |
| **Cell type** | HEK293T | K562 | HEK293T |
| **Feature type** | Peaks formed by chimeric reads | Peaks formed by chimeric reads | Peaks formed by IP reads normalized by input reads |

| **Number of**<br>**replicates (r)** | Two | Four | Four |
| --- | --- | --- | --- |
| **Proxy for truly**<br>**reproducible**<br>**features ($K_i = 1$)** | Chimeric peaks<br>annotated with the<br>transfected miRNA | Chimeric peaks on<br>3' UTR with<br>positive seed<br>matching | Peaks on introns or<br>3' UTR with<br>RBFOX2 motif |

The protocol details for both technologies can be found in Manakov et al. (2022) and Van Nostrand et al. (2016), respectively. Briefly, miR-eCLIP is an extension of the standard eCLIP protocol using an anti-AGO2 antibody and modified to enable chimeric ligation of miRNA and target mRNA. RBFOX2 RBP-eCLIP (RBFOX2-eCLIP), on the other hand, was obtained by applying RBP-eCLIP against RBFOX2, an RNA-binding protein with a widespread role in several cellular mechanisms and tissue-specific effects (Arya *et al*., 2014). RBP-eCLIP extends the single-end eCLIP protocol (Van Nostrand *et al*., 2016) by optimizing several aspects of the original technique, like cell input requirement, UV crosslinking settings, and the lysis protocol.

The resulting sequencing data went through multiple steps of preprocessing and then clusters of aligned reads were found using CLIPper (v2.0.1). For the miR-eCLIP data sets, each cluster was annotated with the names of miRNAs responsible for that target. For the RBFOX2 eCLIP, the IP levels were normalized with input levels to calculate fold enrichments, and p-values were calculated using a Yates’ Chi-Square test (or Fisher Exact Test if the observed or expected read number was below 5). In all datasets, peaks were annotated using transcript information from GENCODE release 41 (GRCh38.p13) (Frankish *et al*., 2022) with the following priority hierarchy to define the final annotation of overlapping features: protein-coding transcript (CDS, UTRs, intron), followed by noncoding transcripts (e.g., exon, intron).

### 2.2 Reproducibility assessment methods: models and algorithms

#### 2.2.1 Estimation of mixture copula model parameters

Suppose 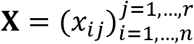 is a matrix with positive values standing for significance or intensity of *n* omic features in r technical replicates. As in standard IDR, gIDR and mIDR, used a copula mixture model on **X**. In the case of mIDR, the model fits in every pairwise combination of replicates. The derivation of the copula model and parameters estimation is as follows. First, feature values are transformed according to the probability integral transform (Savits, 1994) and re-scaled to avoid infinities:

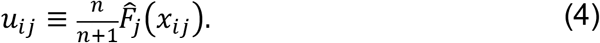

Where 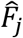 is the empirical cumulative probability distribution (ECDF) of omic feature values across replicate *j*.

Second, the vector of model parameter values is initialized 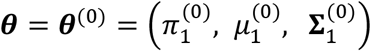. Where 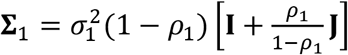, 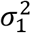 is the reproducible class variance, *ρ*_1_ is the reproducible class correlation, **I** is a rxr matrix with ones in the diagonal and zero otherwise and **J** is rxr matrix with all ones. In the case of the irreproducible class, ∑_0_ = **I**.

Third, **X** is turned into “pseudo data” 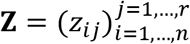, where *z*_*ij*_ = *G*^−1^(*u*_*ij*_|θ), with

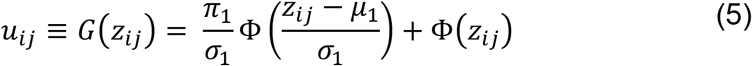

and Φ is the univariate standard normal cumulative density function. The value of *G*^−1^ is obtained by calculating *G* on regular grid of 1,000 *u*_*ij*_ values followed by linear interpolation as in Li et al. (2011).

Then, an EM algorithm on the augmented pseudo data ***y***_*i*_ = (*K*_*i*_, ***z***_*i*_) is used to obtain ***θ***^(*t*)^ (i.e., the estimates of parameter values at iteration *t*) where ***z***_*i*_ = [ *z*_*i*1_, …, *z*_*i*r_]. After setting ***θ*** = ***θ***^(*t*)^, the last two steps are repeated until convergence or a maximum value to *t* (*t*_*MAX*_) are reached.

#### 2.2.2 gIDR

The principal difference between IDR and gIDR is that the latter incorporates information from all replicates by using the following complete log-pseudolikelihood formula:

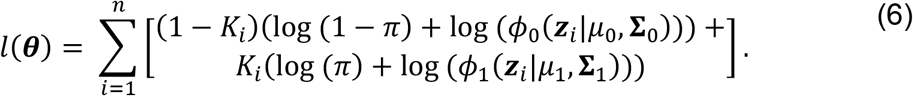

where *ϕ*_*K*_ is the probability density function of a “General” Multivariate Normal distribution, with dimension equal to the number of replicates r (see equation (2))

Since the model is in essence a mixture of two normal distributions on the pseudo data, standard EM formulas apply (Xu *et al*., 2016). Hence, for the Expectation step (*E-step*) we have

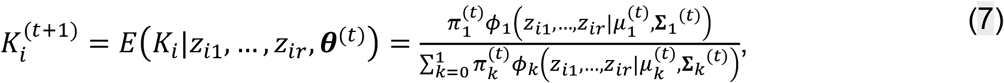

For the Maximization step (*M-step*), we have:

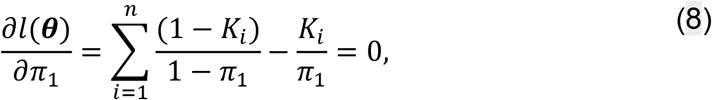

and

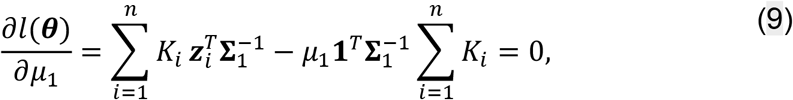

and

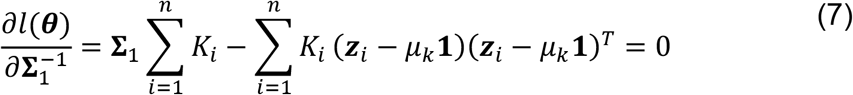

Where in equation (10) we used a) the derivatives of the logarithm of a matrix’s determinant with respect to 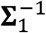 and b) the derivative of the trace of the product between a matrix and 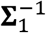 with respect to 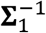 (Dwyer, 1967).

The maximum likelihood estimates of each parameter in the (*t* + 1)-th iteration are given by the following expressions:

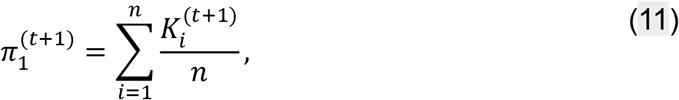

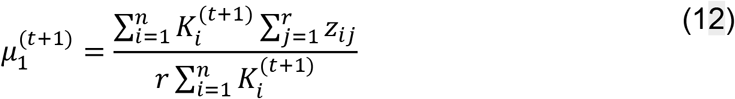

and

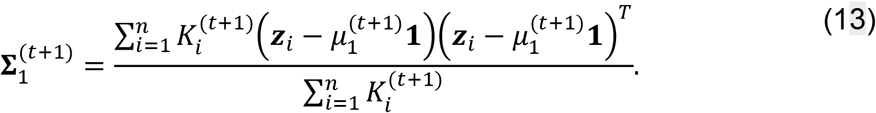

Finally, since 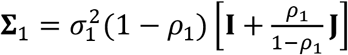, we have:

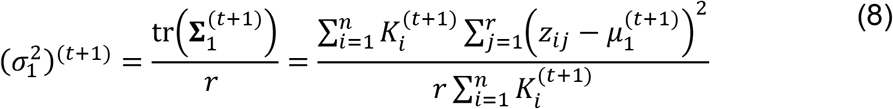

and

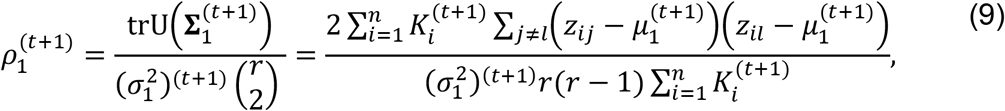

where tr and trU are the trace and upper-triangular trace of a matrix, respectively, and 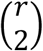 is the number r replicates combinations in two. Finally, the local IDR is approximated as 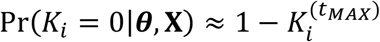, where *t*_*MAX*_ is the number of iterations at convergence. Notice that when r = 2, the *E-step* and *M-step* formulas above reduce to ones derived for standard IDR (see Li *et al*, 2011 and supplementary materials there).

#### 2.2.3 mIDR

Method mIDR is derived using a simple heuristics method that resembles a meta-analysis score (Shelby and Vaske, 2008). First, we perform standard IDR in each one of every pairwise combination of replicates. Then, a meta score is calculated to get the probability of a feature being irreproducible in at least one of the 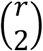 comparisons:

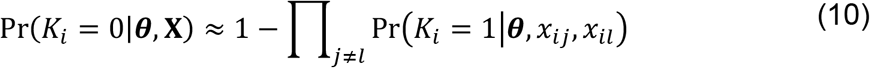

#### 2.2.4 eCV

The eCV method assumes omic feature values across replicates ***y***_*i*_ = [*y*_*i*1_, *y*_*i*2_, …, *y*_*i*r_] are positive and approximately Normal. If *x*_*ij*_ stands for the number of aligned sequencings reads or another measurement of the feature’s intensity, with distribution approximately log-Normal, taking *y*_*ij*_ = log (*x*_*ij*_+∈), where ∈ is an added positive value that prevents infinite values, yields the same distribution as in (1).

The estimated eCV is calculated as:

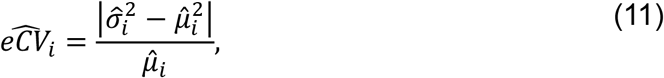

where 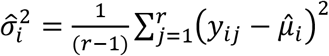 and 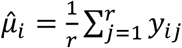. Smaller values of 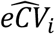 would show reproducibility and vice versa.

To make inferences on the reproducibility of feature *i*, we first compute the following expression

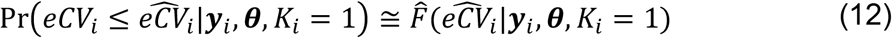

With 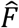 being the ECDF of *eCV*_*i*_. Equation (18) is analogous to a p-value. The larger the value of 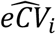, the larger the value of (18).

Second, we apply the Bayes theorem and the chain rule of probability to connect (18) with Pr(*K*_*i*_|***θ*, X**):

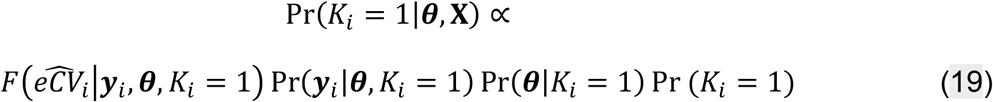

In a fully Bayesian treatment of (19), prior distributions for ***θ***,*K*_*i*_ and their hyperparameters (e.g., *π*) might be used. For eCV, however, we assume a somewhat radical “Empirical Bayes” (EB) (Morris, 1983) method, where we set 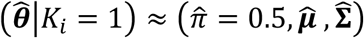. The quantities 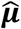 and 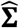 are, respectively, the vector of means and the variance-covariance matrix across features. The values of 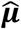 and 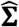 are assumed as good proxies for the parameters of prior distribution of reproducible features.

Third, EB estimates are plugged into equation (1) to randomly sample 10,000 values of 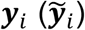. eCV is calculated in every 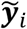 to obtain 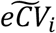 values. The ECDF is used to obtain 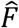. Lastly, 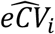 is plugged in 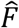 to obtain an estimate of (19).

### 2.3 Performance assessment

#### 2.3.1. Defining a common reproducibility index across methods

Methods were compared using estimates of Pr (*K*_*i*_ = 0|***θ*, X**) as reproducibility index 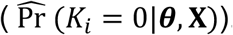. This index can be interpreted as the probability of an omic feature belonging to the irreproducible class. In the case of IDR and its extensions, gIDR and mIDR, the value Pr (*K*_*i*_ = 0|***θ*, X**) was approximated by the local IDR score (Li et. al 2011). For eCV, Pr (*K*_*i*_|***θ*, X**) was approximated using parametric bootstrap.

#### 2.3.1 Performance quantification

In all scenarios, the range of 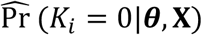 was split into 20 thresholds accommodated in the vector τ = [τ_1_, τ_2_, …, τ_20_]. The thresholds covered the range between 0 and 100%, with each value τ_m_ splitting features into reproducible or not. Each threshold was calculated 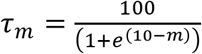, where m is an integer from 1 to 20. The individual values of the true state of each feature *i* (*K*_*i*_) were imposed in each type of scenario. For simulations, the value of *K*_*i*_ was a priori determined in each simulated data set, depending on the value of *π*. For the experimental data sets, the value of *K*_*i*_ was imposed based on biological knowledge from each system (Table 2). In all scenarios, the “true” feature state was used to estimate true positives rate (TPR), showing true reproducible signal, and true negative rates (TNR), showing true irreproducible noise.

#### 2.3.2 Inference and summary of performance values

TPR and TNR across τ values were calculated with the R package *pROC* (Robin *et al*., 2011). For simulation scenarios, performance values were summarized in terms of median and 90% confidence intervals across simulations. Performance values in experimental data were summarized by the median and 90% confidence intervals across 2,000 bootstrap samples.

### 2.4 Software implementation and data availability

All methods and simulation scenarios are implemented as functions of the R package *eCV*. All the software required to generate the results presented here are preinstalled in a publicly available Docker image *ecv_results*. The image can be retrieved via *docker pull ecvpaper2024/ecv_results:latest*. The miR-eCLIP data from transfections are available from the GEO accession <u>GSE198250</u> (miR-1 transfected samples, GSM5942220-1, miR-124 transfected samples GSM5942222-3). The RBFOX2 eCLIP data are available from the GEO accession <u>GSE248884</u> (samples GSM7921694-701). The miR-eCLIP data from wild-type K562 cells are available upon request.

## 3 RESULTS

The four reproducibility assessment methods (**IDR, gIDR, mIDR**, and **eCV**) were used to discriminate between reproducible and irreproducible features. Performance of each method was evaluated in simulated and experimental data sets. In the case standard IDR was used in settings with more than two replicates, only the first two were used. An increasing number of features can significantly increase the computational costs of methods in the IDR family. Although eCV is less affected by the total number of features, we narrowed the analysis to only chromosome one to maintain consistency across comparisons. Therefore, when comparing scenarios, we prioritized performance metrics based on proportions to minimize the effect of a raw number of features. The instructions for generating results using simulated data can be located in Supplementary File S1. Additionally, instructions for generating results from the experimental data for miR-eCLIP transfection, miR-eCLIP wild-type data, and RBFOX2-eCLIP are provided in Supplementary Files S2, S3, and S4, respectively.

### 3.1 Methods performance in simulated scenarios

#### 3.1.1: Simulated Scenario 1

*Many highly correlated, truly reproducible features exist*. This scenario represented simulations where many highly correlated features were reproducible (Figure 1). Performance of the IDR family in this scenario was more robust to varying reproducibility thresholds τ than eCV, in both true positive (TPR) and true negative rates (TNR). The method gIDR was consistently high in both TPR and TNR, particularly for increasing number of replicates r. The performance of mIDR, on the other hand, was more variable, with TNR increasing with r while TPR reaching lower values than the ones achieved with standard IDR. Lastly, the performance of method eCV marginally increased with r but did not reach the TPR performance achieved by IDR and its extensions. Both eCV and mIDR, however, achieved competitive performance at τ’s values between 50-75%

**Figure 1:**
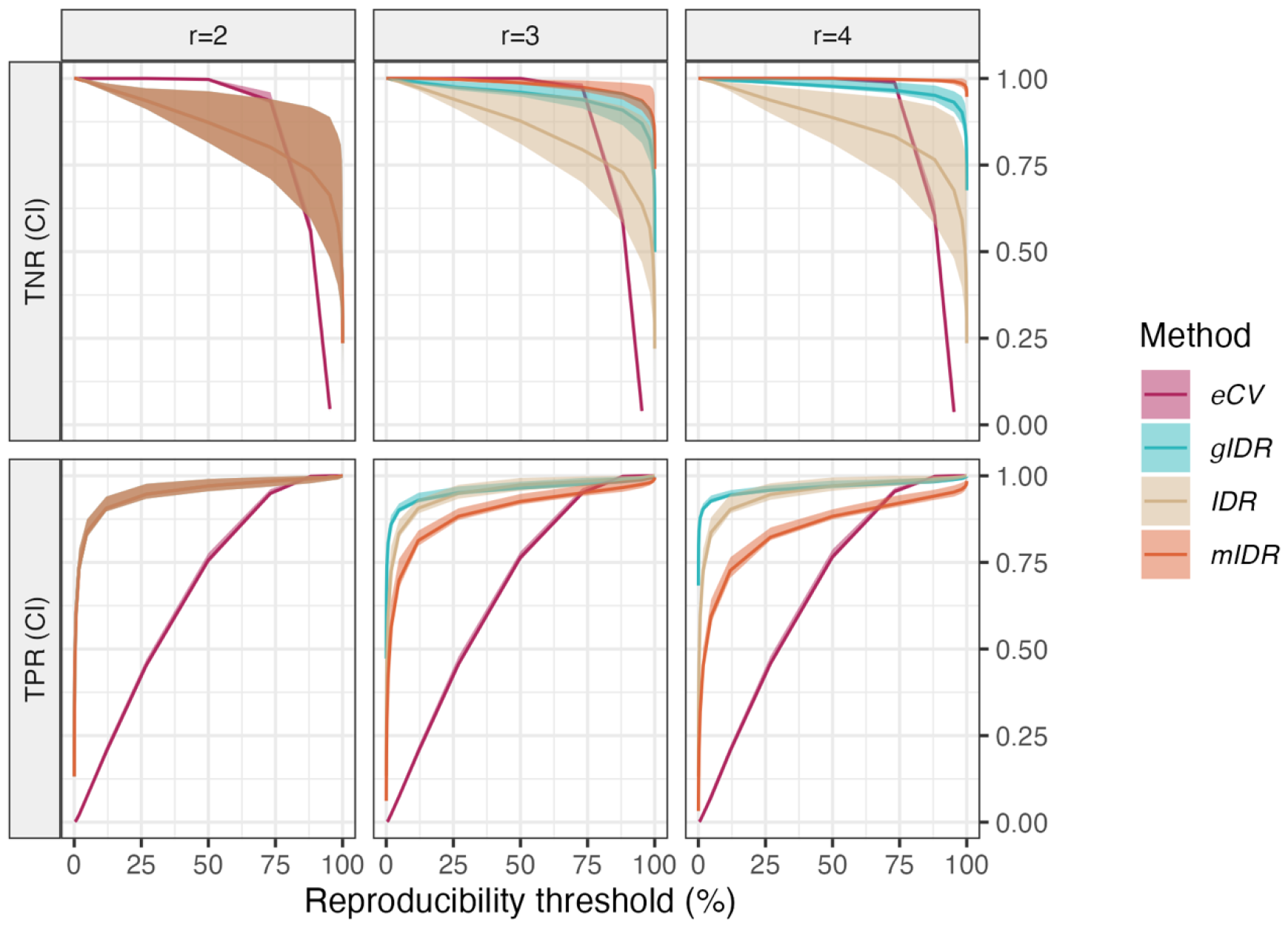
Methods performance in simulated scenario 1. *Many strongly correlated features of genuine signal*. Row panels are median and 90% credible intervals (**CI**) of true negative rate (**TNR**, top) and true positive rate (**TPR**, bottom) rates across 100 simulated data sets. Column panels are varying number of replicates (*r* = 2, 3, or 4). The Y-axis is TPR or TNR values and the X-axis is the threshold on the reproducibility indices.

#### 3.1.2: Simulated Scenario 2

*Few weakly correlated truly reproducible features* This scenario represented situations where most features belonged to the irreproducible class and were lowly correlated (Figure 2). The overall performance values in this scenario were like the ones in scenario 1, with some differences. For example, gIDR gained even more accuracy over IDR (through increases in TNR values), but not power (no significant difference between IDR and gIDR’s TPR values). The performance of eCV was once more overly sensitive to variations in thresholds τ. However, eCV’s performance steadily increased with r, reaching competitive values at τ ≈ 50%. The method mIDR had marginally higher TNR values compared to IDR, but its TPR values significantly decreased with the number of replicates r.

**Figure 2:**
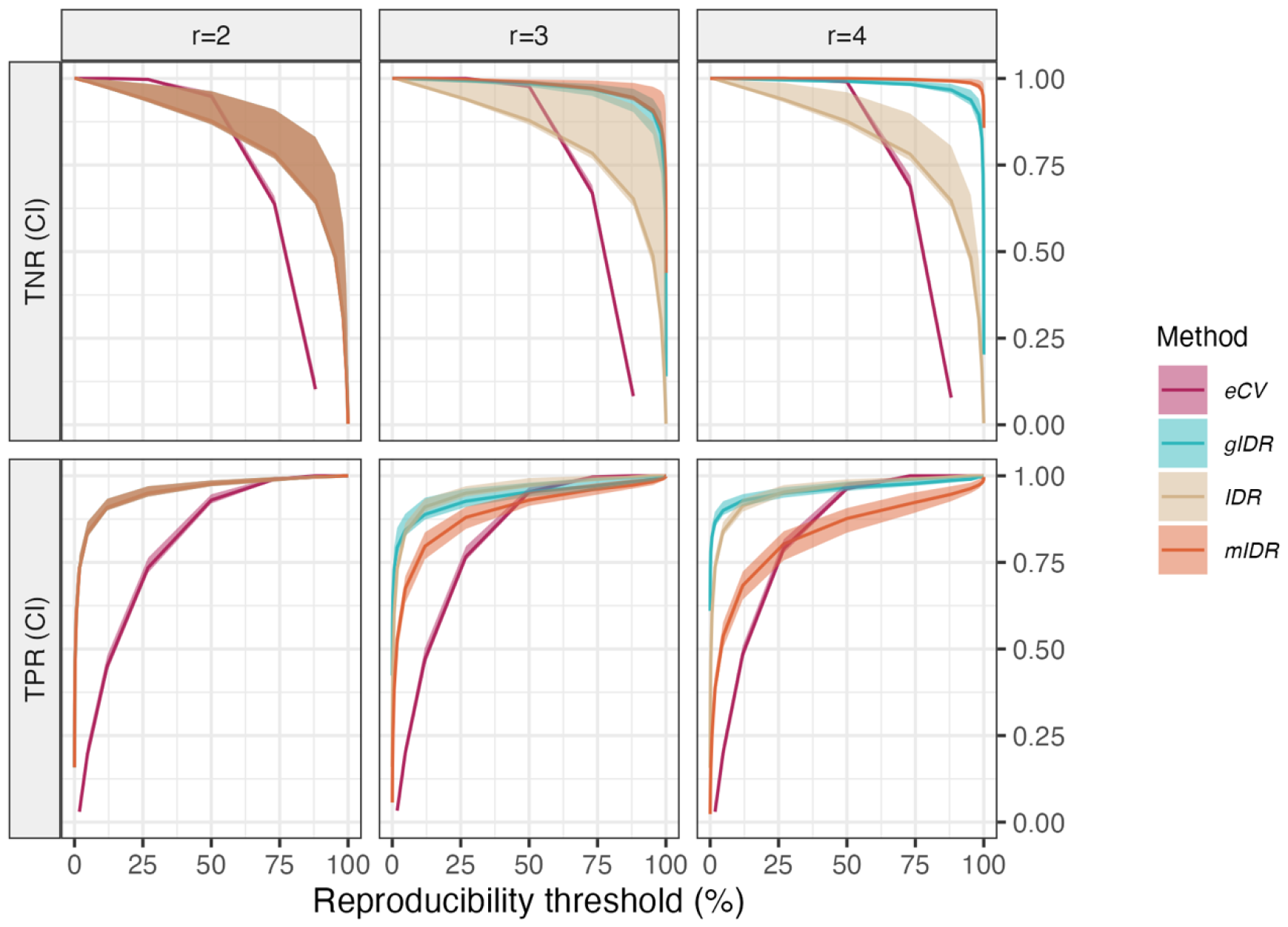
Methods performance in simulated scenario 2. *Few lowly correlated features of genuine signal*. Row panels are median and 90% credible intervals (**CI**) of true negative rate (**TNR**, top) and true positive rate (**TPR**, bottom) rates across 100 simulated data sets. Column panels stand for varying number of replicates (*r* = 2, 3, or 4). The Y-axis is TPR or TNR values and the X-axis is the threshold on the reproducibility indices.

#### 3.1.3: Simulated Scenario 3 Few and highly correlated truly reproducible features

The final simulated scenario represented situations with an even smaller number of strongly correlated reproducible features (Figure 3). In this scenario, eCV had the best overall performance across number of replicates *r*, achieving high TPR and TNR even for smaller thresholds τ.. Contrary to the earlier simulated scenarios, the performance of gIDR became severely affected by the increasing number of replicates *r*. Method mIDR, on the other hand, systematically improved with *r*, achieving the best TNR values at *r* = 4.

**Figure 3:**
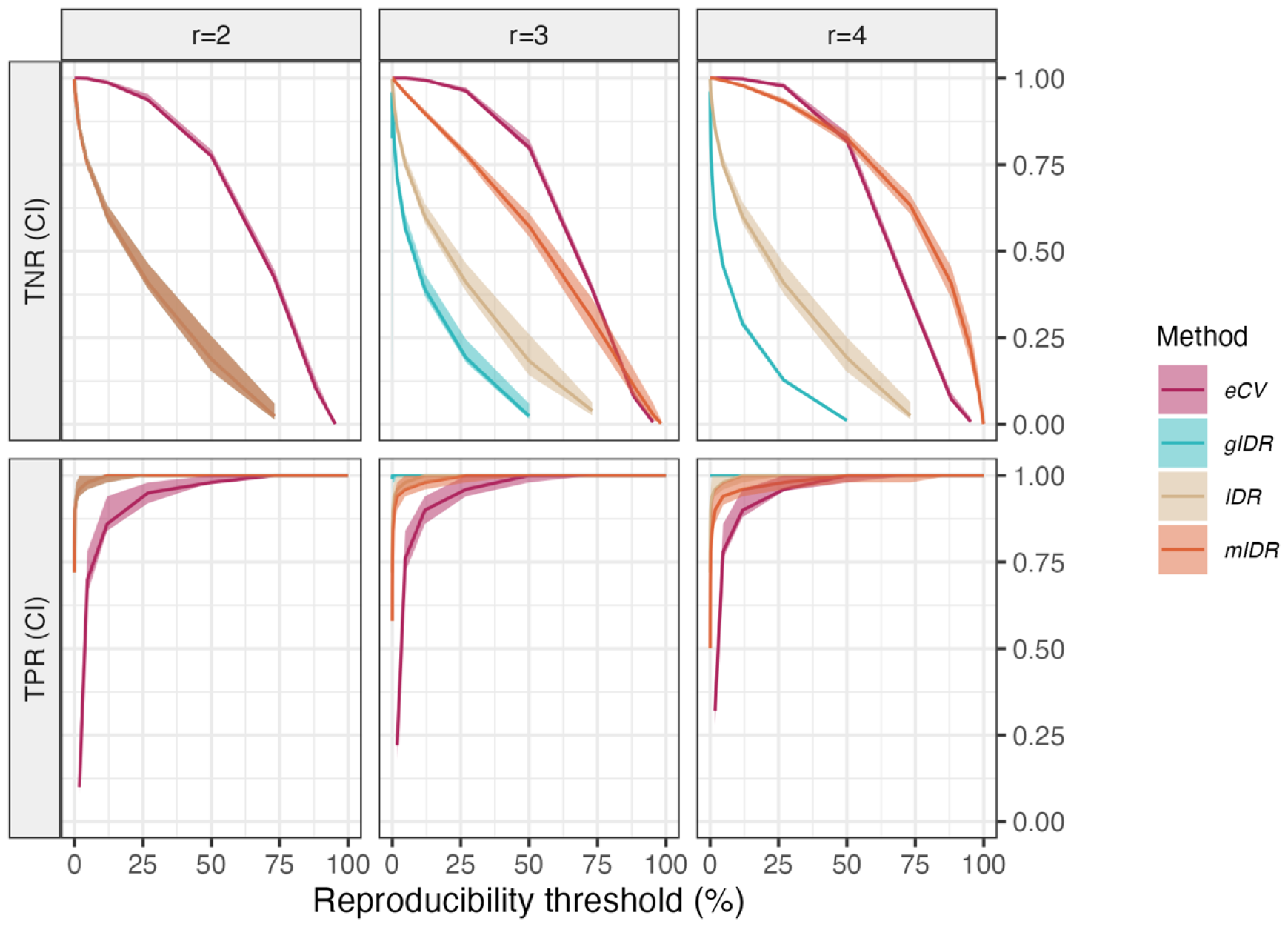
Methods performance in simulated scenario 3. *Few strongly correlated features of genuine signal*. Row panels stand for median and 90% credible intervals (**CI**) of true negative rate (**TNR**, top) and true positive rate (**TPR**, bottom) rates across 100 simulated data sets. Column panels are varying number of replicates (*r* = 2, 3, or 4). The Y-axis is TPR or TNR values and the X-axis is the threshold on the reproducibility indices.

**Figure 4:**
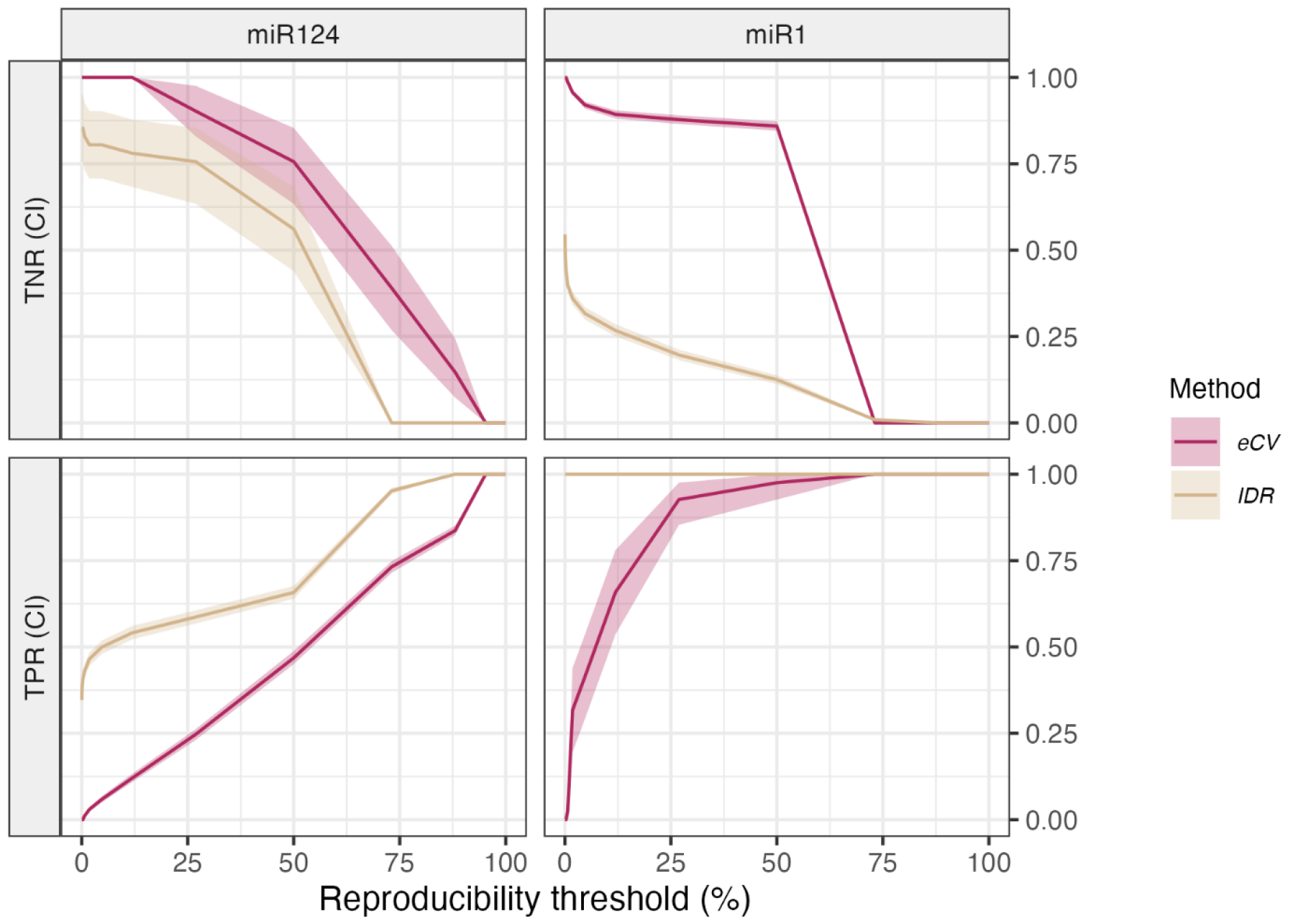
Methods performance in experimental scenario 1: *miR-eCLIP data from transfected cells*. Row panels are median and 90% credible intervals (**CI**) of true negative rate (**TNR**, top) and true positive rate (**TPR**, bottom) rates. Column panels are the transfections: *hsa-miR-124-3p* (**miR124**, left) and *hsa-miR-1-3p* (**miR1**, right). The Y-axis is TPR or TNR values and the X-axis is the threshold on the reproducibility indices.

**Figure 5:**
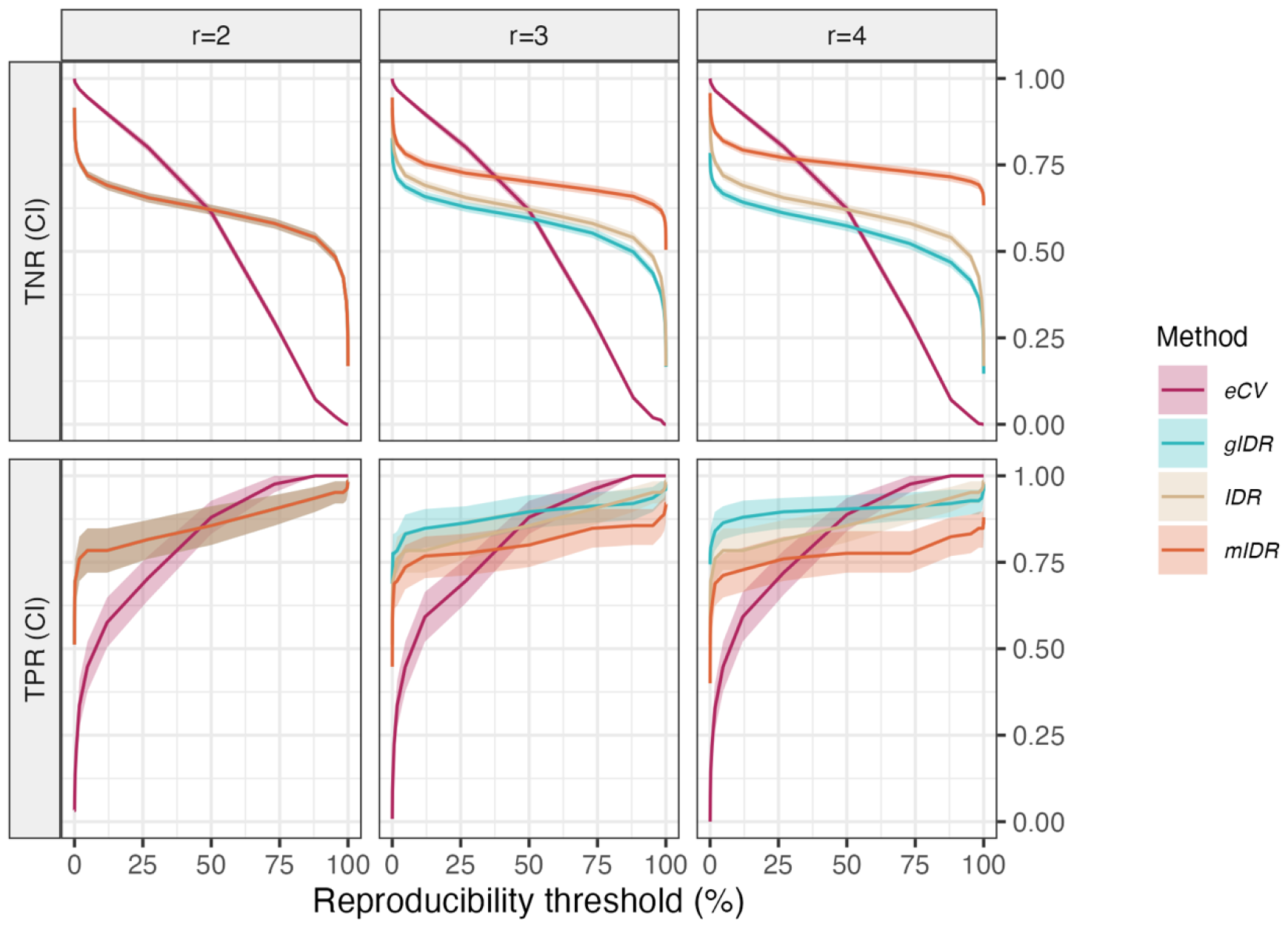
Methods performance in experimental scenario 2: *miR-eCLIP data from wild-type cells*. Row panels are median and 90% credible intervals (**CI**) of true negative rate (**TNR**, top) and true positive rate (**TPR**, bottom) rates. Column panels are varying number of replicates (*r* = 2, 3, or 4). The Y-axis is TPR or TNR values and the X-axis is the threshold on the reproducibility indices.

**Figure 6:**
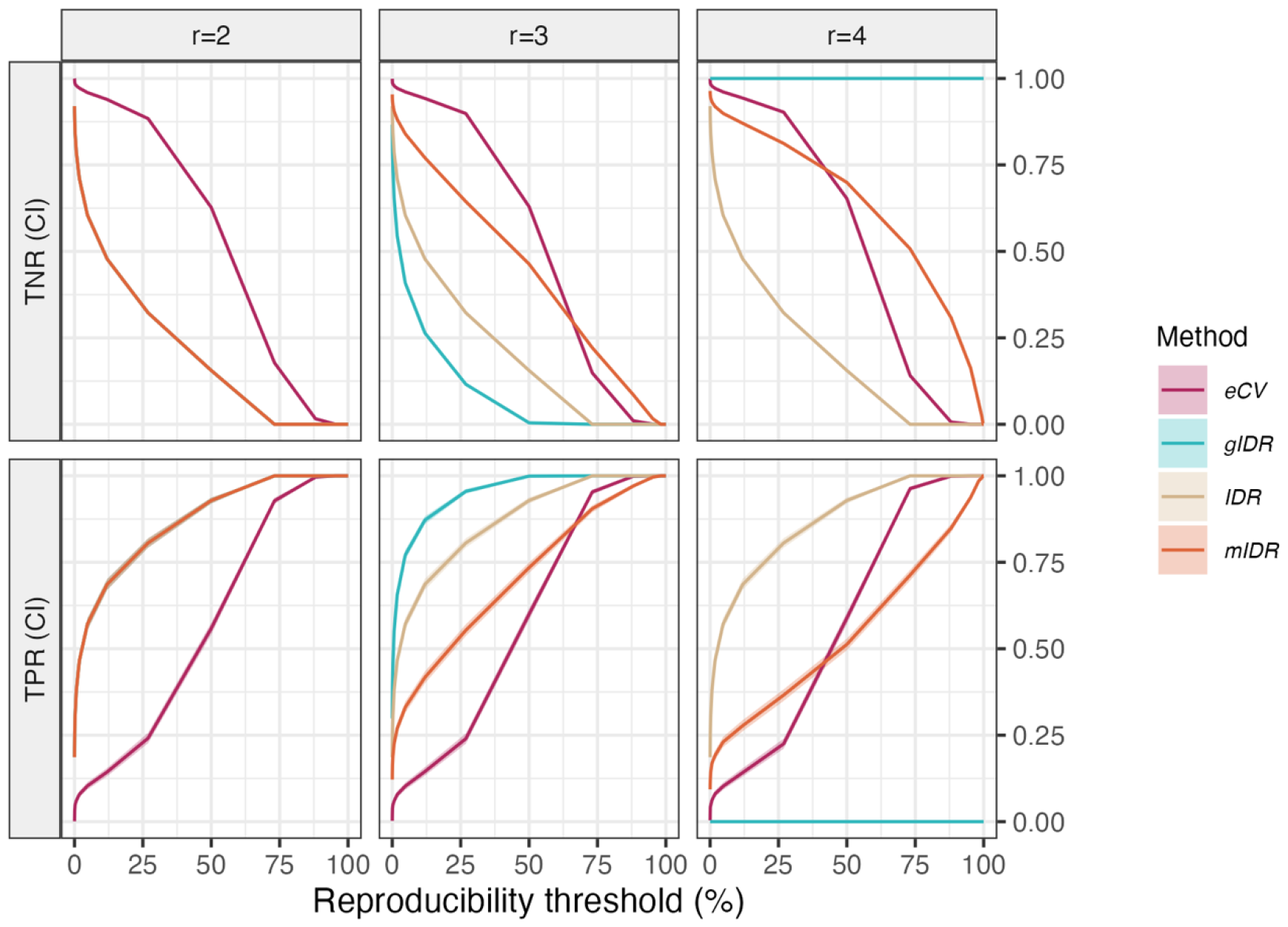
Methods performance in experimental scenario 3. *RBFOX2 eCLIP data from wildtype cells*. Row panels are median and 90% credible intervals (**CI**) of true negative rate (**TNR**, top) and true positive rate (**TPR**, bottom) rates. Column panels are varying number of replicates (*r* = 2, 3, or 4). The Y-axis is TPR or TNR values and the X-axis is the threshold on the reproducibility indices.

### 3.2 Methods performance in experimental data

#### 3.2.1: Experimental Scenario 1: miR-eCLIP data on transfected cells

The first experimental data set came from chimeric peaks data (peaks formed by chimeric reads coming from mRNA:miRNA interactions) obtained with miR-eCLIP on HEK293T cells. Peaks annotated with any of the two transfected miRNAs: *hsa-miR-124-3p* (miR-124) and *hsa-miR-1-3p* (miR-1), were used as proxies of truly reproducible features. Endogenous mRNA:miRNA chimeras, categorized as irreproducible, were considered as “true negatives.” This selection was justified by the observation that the majority of endogenous chimeras in this dataset exhibited significantly lower intensities compared to those involving transfected miRNAs. Only *n* 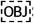 = 3,145). Of these, 1,560 were annotated with miR-124, while only 257 were annotated with miR-1. Confidence intervals at the 95% level suggested a high correlation of the peak intensities for both transfected miRNAs: [0.79, 0.83] for miR-124 and [0.55, 0.70] for miR-1. Since only two technical replicates were available in each transfection, only IDR and eCV were compared.

The results of the method comparisons in the miR-124 transfection resembled those obtained in simulated scenario 1 (i.e., many strongly correlated truly signal features). The results of the methods comparisons in the miR-1 transfection, on the other hand, resembled the ones in scenario 3 (i.e., small number of strongly correlated truly signal features). Although the global behavior of both IDR and eCV was like that in the simulations, eCV had significantly higher global performance than IDR, particularly for **τ** ≈ 50%.

#### 3.2.2: Experimental Scenario 2: miR-eCLIP data from wild-type cells

The second experimental data set also came from chimeric peaks data obtained with miR-eCLIP, but this time from two, three, and four replicates from wild-type K562 cells. Only chromosome 1 peaks were used (*n* = 1,858). Contrary to the earlier scenario, this one consisted of only endogenous chimeras. Therefore, only significantly enriched peaks with positive seed matches on the 3’ UTR were used as proxies of “truly reproducible” features. Although the peaks being “truth” were strongly correlated within groups of genes targeted by 13 miRNA species involved in those peaks (∼0.9), the correlations between groups miRNA significantly dropped.

Results resembled the ones in simulated scenario 2 (i.e., few lowly correlated truly reproducible features). Like in scenario 2, gIDR’s TNR values improved with increasing number of replicates *r*, while TPR values were not significantly different than IDR’s. Also, in agreement with simulated scenario 2, eCV varied more quickly with thresholds τ than the other methods, while also steadily improving with the number of replicates. As before, eCV matched the performance of every other method when **τ** ≈ 50%, except for mIDR’s TNR. The performance of mIDR also mirrored simulated scenario 2, except that here, its TNR values were significantly higher than any other method.

#### 3.2.3: Experimental Scenario 3: RBFOX2 eCLIP data on wildtype cells

The third and last experimental data set came from peak data derived from an RBFOX2 eCLIP on HEK293T cells for 2, 3, and 4 replicates. Peaks enriched with the RBFOX2 binding motif and mapping onto the 3’ UTR were proxies for “truly reproducible” features. Only chromosome 1 peaks were used (*n* = 24,703). Of these, only 5,173 fulfilled both conditions, with approximately 20% of the peaks having the RBFOX2 motif and approximately 13% mapping onto the 3’ UTR. Correlations among these peaks were also highly significant: confidence intervals at 95% = [0.60, 0.63].

Results in this scenario resembled the ones for simulated scenario 3, distinguished by only a handful of highly correlated truly reproducible features. As for simulated scenario 3, eCV reached higher TPR values with increasing r while systematically keeping higher TNR, particularly at τ around 50%. Similarly, mIDR improved its TNR values for increasing r. However, contrary to simulation results, its TPR seemed to decrease. Also, consistently with simulated scenario 3, gIDR performance was severely affected, particularly for increasing *r*.

## 4 DISCUSSION

The escalating capacity for multiplexing and diminishing sequencing costs make it possible to work with increasing sample sizes, thereby augmenting the quality of highthroughput data by raising power and precision (Chen et al., 2012; Li et al., 2017). Nevertheless, reproducibility analysis is still needed to ensure valid results. This becomes a non-trivial problem when using standard techniques, such as irreproducible discovery rate (IDR), on more than two replicates (Yang et al., 2014). We propose three novel methods to overcome this limitation as alternatives to the traditional IDR. Results from both simulations and experimental data suggest that the methods presented here can outperform IDR in several relevant biological scenarios.

Both simulated and experimental data scenarios were chosen to show some semblance to each other. For example, the chimeric peaks in experimental scenario 1 (cells transfected with miR-124 or miR-1) annotated with the respective transfected miRNA resulted in significantly high correlations. These high correlations and the fact that miR-124 had more than 83% of annotated peaks, while miR-1 had about 14%, could explain the strong resemblance between this data set and simulation scenarios 1 and 3, standing for many and few strongly correlated truly reproducible features, respectively. A similar explanation can be applied to the behaviors between simulated scenarios two and three and the second miR-eCLIP and RBFOX2 eCLIP data sets.

Of all the methods compared, the general IDR (gIDR) was particularly powerful at detecting truly reproducible features while discarding truly irreproducible ones in many of the scenarios tested. This behavior manifested in two opposite situations: many highly correlated reproducible features (e.g., simulation scenario 1) and few lowly-correlated reproducible features (e.g., simulation scenario 2). Unfortunately, the performance of gIDR significantly dropped when even fewer but highly correlated reproducible features were present (e.g., simulation scenario 3). In those cases, gIDR underperformed with an increasing number of replicates. One compelling explanation for these behaviors is the high sensitivity of multivariate normal components to the misspecification of initial parameter values in combination with ill-conditioned variance-covariance matrices (Gray, 1994; Lo, 2011). Consider for example the determinant of the variance-covariance matrix of the reproducibility component 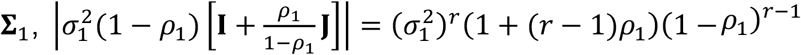, where as before, 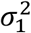 and *ρ*_1_ the reproducible class variance and correlation. If, say 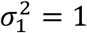 and *ρ*_1_ = 0.9, then the determinant 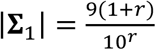 quickly approaches zero as r grows, making **∑**_1_ near singular. This, combined with near-boundary values of the mixing parameter *π* (e.g., 0.05, as in simulated scenario 3), might artificially push features into one mixture components, regardless of the actual reproducibility state. Potential solutions to these issues are application of regularization during estimations of **∑**_1_ (Liu and Palomar, 2019) and preoptimized initial values of parameters parameter values (Panić *et al*., 2020). Meta IDR (mIDR) performance varied more than gIDR, being particularly high in simulated scenario 3. Although the experimental RBFOX2 eCLIP scenario resembled simulated scenario 3, the method did not perform as well, particularly in terms of TPR. Nevertheless, it seems that mIDR would increase its performance with an increasing number of replicates and be particularly good in situations where only a few features are truly reproducible. Although the performance of mIDR would increase with r, since traditional IDR is applied in every pairwise combination, the number of total comparisons would increase quadratically. Although the calculation of IDR in every pair of replicates can be achieved in parallel, it is also worth mentioning potentially high computational costs if several replicates are jointly analyzed.

The global performance of eCV was variable across scenarios. Although with the highest sensitivity to the choice of the threshold on the reproducibility values, eCV was undoubtedly the more consistent method across varying numbers of replicates. eCV was also less potent than IDR when most features were reproducible and highly correlated. These facts point to eCV being a more stringent method than IDR, potentially due to the heavy prior information integrated into the model via Empirical Bayes. Nevertheless, eCV still detected the most reproducible features (e.g., those with higher feature intensities, Supplementary Figure S1) while calling fewer false positives. In addition, eCV also had significantly more global accuracy, most clear in the miR-124 miR-eCLIP experiment. As opposed to gIDR, eCV would also have higher performance in scenarios with fewer reproducible features.

Further research on the applicability of a fully Bayesian treatment of eCV could alleviate the potential stringency of the method. Nevertheless, due to the low sample sizes typically used in high-throughput experiments, a strong prior via EB, like the one used in the *limma* suit of models (Ritchie et al., 2015) could augment power. The ability to detect highly reproducible features, combined with its high TNR across different scenarios, makes eCV a competitive method, particularly when a threshold of 50% was used. This threshold is also compelling since it matches the intuition of calling features as reproducible; their values surpass what is expected from entirely random behavior.

All the methods were compared on the same scale. In the case of IDR and its extensions, this reduced to working with the so-called local IDR. In the case of eCV, this involved using the probability of the estimated eCV being significantly larger than values expected for reproducible features. Although a method for FDR control via eCV was not tested, a similar approach to the one used by IDR could be implemented. This would involve plugging the eCV’s reproducibility indices into a multiple comparison approach, like the one adopted by IDR following Sun and Cai (2007).

We have shown that multivariate alternatives to standard IDR might perform highly, but choosing the best model still is a non-trivial issue. Although the results shown here supply some guidelines, due to most biological problems not having a clear sign of “truth,” deciding when to use one or another method is not trivial. Therefore, deciding which method would perform the best would depend on various aspects, such as what type of statistical error one wishes to control the most. Due to its robustness to varying sample sizes, stable behavior, and high accuracy, we favor eCV as a suitable high-performance method across many contrasting scenarios.

## Conflict of interest

Eclipsebio funded and conducted the research presented in this paper. The authors are paid employees of Eclipsebio but have no financial interests or personal relationships that could potentially bias the results or interpretation of the data.

## Notes

https://github.com/eclipsebio/eCV

### Summary of Updates

Change corresponding author

## REFERENCES

Arya, A.D. et al. (2014) RBFOX2 protein domains and cellular activities. Biochem. Soc. Trans., 42, 1180–1183.

Chen, Y. et al. (2011) MM-ChIP enables integrative analysis of cross-platform and between-laboratory ChIP-chip or ChIP-seq data. Genome Biol., 12, R11.

Chen, Y. et al. (2012) Systematic evaluation of factors influencing ChIP-seq fidelity. Nat. Methods, 9, 609–614.

Dwyer, P.S. (1967) Some Applications of Matrix Derivatives in Multivariate Analysis. J. Am. Stat. Assoc., 62, 607–625.

Frankish, A. et al. (2022) GENCODE: reference annotation for the human and mouse genomes in 2023. Nucleic Acids Res., 51, D942–D949.

Genest, C. and Nešlehová, J. (2013) Copulas and Copula Models. In, Encyclopedia of Environmetrics. John Wiley & Sons, Ltd.

Gray, G. (1994) Bias in misspecified mixtures. Biometrics, 50, 457–470.

Gu, S. et al. (2009) The biological basis for microRNA target restriction to the 3’ untranslated region in mammalian mRNAs. Nat. Struct. Mol. Biol., 16, 144–150.

Haecker, I. et al. (2012) Ago HITS-CLIP Expands Understanding of Kaposi’s Sarcoma-associated Herpesvirus miRNA Function in Primary Effusion Lymphomas. PLOS Pathog., 8, e1002884.

Heinz, S. et al. (2010) Simple combinations of lineage-determining transcription factors prime cis-regulatory elements required for macrophage and B cell identities. Mol. Cell, 38, 576–589.

Landt, S.G. et al. (2012) ChIP-seq guidelines and practices of the ENCODE and modENCODE consortia. Genome Res., 22, 1813–1831.

Li, C.-I. et al. (2017) Power and sample size calculations for high-throughput sequencingbased experiments. Brief. Bioinform., 19, 1247–1255.

Li, Q. et al. (2011) Measuring reproducibility of high-throughput experiments. Ann. Appl. Stat., 5.

Liu, J. and Palomar, D.P. (2019) Regularized robust estimation of mean and covariance matrix for incomplete data. Signal Process., 165, 278–291.

Lo, Y. (2011) Bias from misspecification of the component variances in a normal mixture. Comput. Stat. Data Anal., 55, 2739–2747.

Manakov, S.A. et al. (2022) Scalable and deep profiling of mRNA targets for individual microRNAs with chimeric eCLIP. 2022.02.13.480296.

Morris, C.N. (1983) Parametric Empirical Bayes Inference: Theory and Applications. J. Am. Stat. Assoc.

Mundade, R. et al. (2014) Role of ChIP-seq in the discovery of transcription factor binding sites, differential gene regulation mechanism, epigenetic marks and beyond. Cell Cycle, 13, 2847–2852.

Nicholson, G. and Holmes, C. (2017) A note on statistical repeatability and study design for high-throughput assays. Stat. Med., 36, 790–798.

Panić, B. et al. (2020) Improved Initialization of the EM Algorithm for Mixture Model Parameter Estimation. Mathematics, 8, 373.

Revilla-i-Domingo, R. et al. (2012) The B-cell identity factor Pax5 regulates distinct transcriptional programmes in early and late B lymphopoiesis. EMBO J., 31, 3130–3146.

Robin, X. et al. (2011) pROC: an open-source package for R and S+ to analyze and compare ROC curves. BMC Bioinformatics, 12, 77.

Rozowsky, J. et al. (2009) PeakSeq enables systematic scoring of ChIP-seq experiments relative to controls. Nat. Biotechnol., 27, 66–75.

Savits, T.H. (1994) On integration, substitution and the probability integral transform. Stat. Probab. Lett., 21, 173–179.

Shelby, L.B. and Vaske, J.J. (2008) Understanding Meta-Analysis: A Review of the Methodological Literature. Leis. Sci., 30, 96–110.

Soutschek, M. et al. (2022) scanMiR: a biochemically based toolkit for versatile and efficient microRNA target prediction. Bioinformatics, 38, 2466–2473.

Sun, W. and Cai, T.T. (2007) Oracle and Adaptive Compound Decision Rules for False Discovery Rate Control. J. Am. Stat. Assoc., 102, 901–912.

Tsukahara, H. (2005) Semiparametric estimation in copula models. Can. J. Stat., 33, 357–375.

Van Nostrand, E.L. et al. (2016) Robust transcriptome-wide discovery of RNA-binding protein binding sites with enhanced CLIP (eCLIP). Nat. Methods, 13, 508–514.

Xu, J. et al. (2016) Global Analysis of Expectation Maximization for Mixtures of Two Gaussians. In, Advances in Neural Information Processing Systems. Curran Associates, Inc.

