## Supplementary File S1 for "eCV: Enhanced coefficient of variation and IDR extensions for reproducibility assessment of high-throughput experiments with multiple replicates"

### Simulated data

This document contains the required steps to generate the results for section **3.1 Methods performance in simulated scenarios**. Please follow the instructions in materials and methods within the main manuscript to download and process the experimental data. Pull a Docker image with all the required packages installed (`docker pull ecvpaper2024/ecv_results`). # Data preparation.

Load required packages.

```
library("eCV")
Loading required package: idr
Loading required package: mvtnorm
Loading required package: future
Loading required package: future.apply
Loading required package: MatrixGenerics
Loading required package: matrixStats

Attaching package: 'MatrixGenerics'
The following objects are masked from 'package:matrixStats':

    colAlls, colAnyNAs, colAnys, colAvgsPerRowSet, colCollapse,
    colCounts, colCummaxs, colCummins, colCumprods, colCumsums,
    colDiffs, colIQRDiffs, colIQRs, colLogSumExps, colMadDiffs,
    colMads, colMaxs, colMeans2, colMedians, colMins, colOrderStats,
    colProds, colQuantiles, colRanges, colRanks, colSdDiffs, colSds,
    colSums2, colTabulates, colVarDiffs, colVars, colWeightedMads,
    colWeightedMeans, colWeightedMedians, colWeightedSds,
    colWeightedVars, rowAlls, rowAnyNAs, rowAnys, rowAvgsPerColSet,
    rowCollapse, rowCounts, rowCummaxs, rowCummins, rowCumprods,
    rowCumsums, rowDiffs, rowIQRDiffs, rowIQRs, rowLogSumExps,
    rowMadDiffs, rowMads, rowMaxs, rowMeans2, rowMedians, rowMins,
    rowOrderStats, rowProds, rowQuantiles, rowRanges, rowRanks,
    rowSdDiffs, rowSds, rowSums2, rowTabulates, rowVarDiffs, rowVars,
    rowWeightedMads, rowWeightedMeans, rowWeightedMedians,
    rowWeightedSds, rowWeightedVars



 __________________________________________________________________
|                        ______     __                         |
|                 ___   / ___\ \   / /                         |
|                / _ \ | |    \ \ / /                          |
|               |  __/ | |___  \ V /                           |
|                \___|  \____|  \_/                            |
|__________________________________________________________________|

        Enhanced Coefficient of Variation and IDR Extensions
                  for Reproducibility Assessment
```

```
This package provides extensions and alternative methods to  IDR  to
measure the reproducibility of omic data with an arbitrary number of
replicates. It introduces an enhanced Coefficient of Variation (eCV)
metric to assess the likelihood of omic features being reproducible.
require("tidyverse")
Loading required package: tidyverse
-- Attaching packages --------------------------------------- tidyverse 1.3.2 --
v ggplot2 3.4.3     v purrr   1.0.2
v tibble  3.2.1     v dplyr   1.1.2
v tidyr   1.3.0     v stringr 1.5.0
v readr   2.1.2     v forcats 0.5.1
-- Conflicts ------------------------------------------ tidyverse_conflicts() --
x dplyr::count()  masks matrixStats::count()
x dplyr::filter() masks stats::filter()
x dplyr::lag()    masks stats::lag()
library("pROC")
Type 'citation("pROC")' for a citation.

Attaching package: 'pROC'

The following objects are masked from 'package:stats':

    cov, smooth, var
library("future")
library("future.apply")
library("reshape")

Attaching package: 'reshape'

The following object is masked from 'package:dplyr':

    rename

The following objects are masked from 'package:tidyr':

    expand, smiths
library("idr2d")
```

Set color palette.

```
res_colors <-
  c(IDR="tan",
    gIDR="#30B7BC",  # bright teal
eCV="#AF275F",  # light magenta
mIDR="#DE653A") #bright orange
```

### Simulation study.

Run simulations. The three simulation scenarios were created with function `eCV::simulate_data`, following Table 1 from Li et al. (2011).

Create a table with all combinations of simulation parameters.

```r
sim_settings <- expand.grid(n_rep = c(4,3,2),
                            scenario=c(1,2,3),
                            method=c("gIDR","mIDR","IDR","eCV"),
                            stringsAsFactors = FALSE) %>%
  mutate(par_com = seq_along(n_rep))
```

Set parameters for each model.

```r
# Set parameters for each model.
methods_params <- list(
  eCV = list(max.ite = 1e4),
  IDR = list(
     mu = 2,
     sigma = 1,
     rho = 0.85,
     p = 0.5,
    eps = 1e-3,
    max.ite = 50
  ),
  mIDR = list(
     mu = 2,
     sigma = 1,
     rho = 0.85,
     p = 0.5,
    eps = 1e-3,
    max.ite = 50
  ),
  gIDR = list(
     mu = 2,
     sigma = 1,
     rho = 0.85,
     p = 0.5,
    eps = 1e-3,
    max.ite = 50
  )
)
```

```r
# Set parallel scheduler.
future::plan(multisession, workers = 4)
perf_res <- NULL
for (i in sim_settings$par_com) {
  set.seed(42)
  cat("i ",i,"\n")
  n_rep <- sim_settings$n_rep[i]
  scenario <- sim_settings$scenario[i]
  method <- sim_settings$method[i]

  tmp <- future_lapply(seq_len(100), function(sim, n_rep, scenario, method) {
    cat("sim = ",sim,"\n")

    sim_data <- simulate_data(n_reps = n_rep, scenario = scenario,
                              n_features = 1000)
```

```
    if(method != "eCV") {
       X <- preprocess(sim_data$sim_data,
                       value_transformation = "identity",
                       jitter_factor=1e-4)
     } else X <- sim_data$sim_data


    rep_index <- mrep_assessment(
         x = X,
         method = method,
         param = methods_params[[method]],
         n_threads = 1
       )$rep_index

   perf <- roc(sim_data$sim_params$feature_group == 2, rep_index, quiet = TRUE)
   perf_thr <- coords(perf,
                        x =  1/(1 + exp(-c(1:20) + 10)),
                        ret=c("threshold","tpr","tnr"))
   perf_thr <- rbind(perf_thr$tpr %>%
   as.data.frame() %>%
   mutate(threshold=1/(1 + exp(-c(1:20) + 10)),
          perf="TPR (CI)") ,
       perf_thr$tnr %>%
   as.data.frame() %>%
   mutate(threshold=1/(1 + exp(-c(1:20) + 10)),
          perf="TNR (CI)"))

   perf_thr$n_rep <- n_rep
   perf_thr$method <- method
   perf_thr$scenario <- scenario
   perf_thr$sim <- sim
   return(perf_thr)
  },n_rep = n_rep,
  scenario = scenario,
  method=method,
  future.seed = NULL) %>% do.call(what = rbind)

  perf_res <- rbind(perf_res, tmp)
}
future::plan(sequential)

# Save results.
colnames(perf_res)[1] <- "value"
saveRDS(perf_res,file="perf_resSim.rds")
```

Arrange results for figure creation.

```
perf_plot_df <-
  perf_res %>%
  group_by(threshold,perf,method,n_rep,scenario)  %>%
   summarise(`50%` = median(value, na.rm = TRUE),
            `5%` = quantile(value, na.rm = TRUE, probs=0.25),
            `95%`= quantile(value, na.rm = TRUE, probs=0.95)) %>%
```

```
  mutate(
         n_rep = factor(n_rep,
                        levels = 2:4,
                        labels = paste0("r=",2:4)) %>% as.character())
`summarise()` has grouped output by 'threshold', 'perf', 'method', 'n_rep'. You
can override using the `.groups` argument.
```

#### Scenario 1.

```
p <- perf_plot_df %>%
  filter(scenario==1) %>%
  ggplot(aes(x=threshold,y=`50%`,color=method)) +
  facet_grid(perf ~  n_rep,switch ="y") +
  geom_line() +
   geom_ribbon(color = NA,
    aes(x = threshold, ymin = `5%`, ymax = `95%`,fill=method),
     alpha = 0.5)  +
  scale_x_continuous(breaks=c(0,0.25,0.5,0.75,1),
                     labels = c("0","25","50","75","100"))+
  scale_color_manual(values=alpha(res_colors,1)) +
  scale_fill_manual(values=alpha(res_colors,0.6)) +
    scale_y_continuous(position = "right")+
   theme_bw() + theme(
                     legend.text = element_text(face = "italic"),
                     strip.background = element_rect(fill=alpha("gray",0.25)),
                  legend.background = element_rect(fill = alpha("white",0.1))) +
   guides(color=guide_legend(ncol=1, override.aes = list(size = 2))) +
   labs(y="",
        fill="Method",
       color="Method",
       x="Reproducibility threshold (%)")
p
```

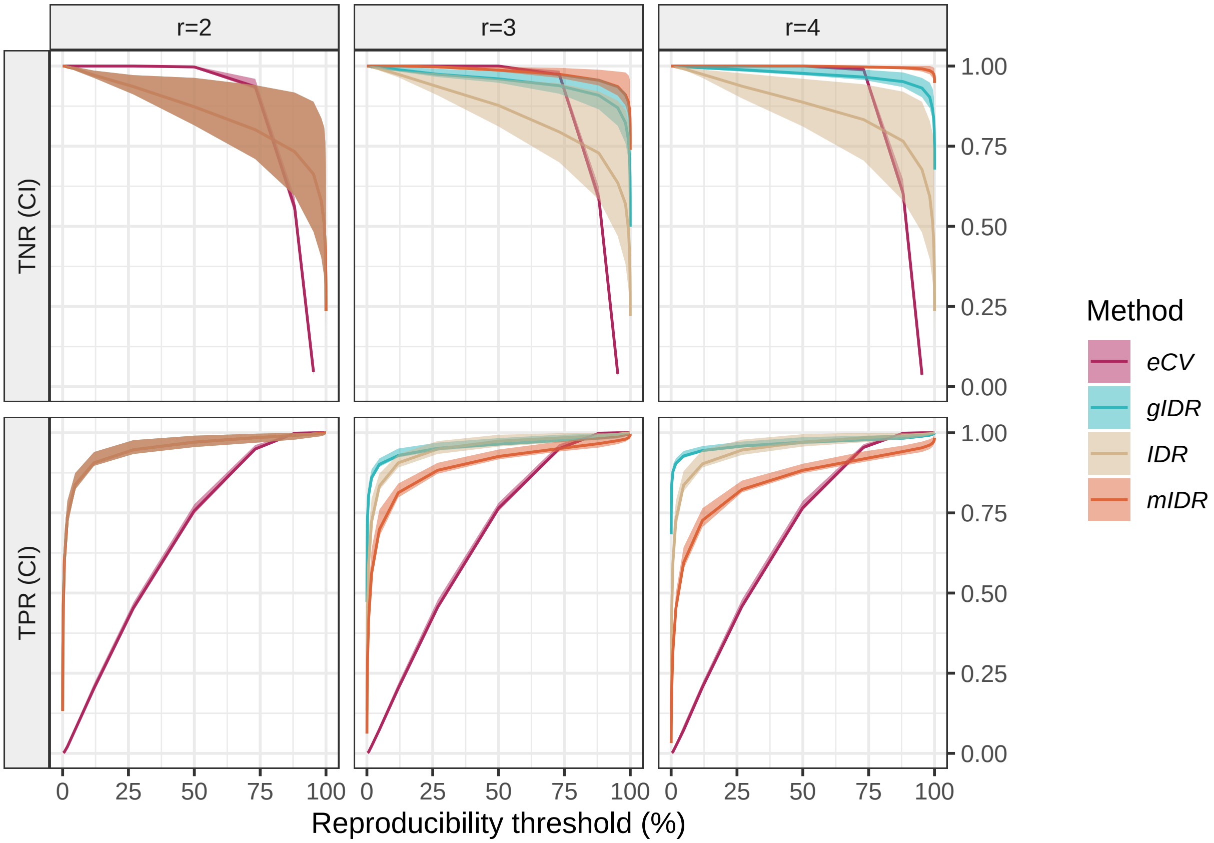


```
ggsave(filename = "Figure1.tiff",plot = p,device = "tiff",dpi=300,
       units = "in",width = 7,height = 5,scale = 0.85)
```

#### Scenario 2.

```
p <- perf_plot_df %>%
  filter(scenario==2) %>%
  ggplot(aes(x=threshold,y=`50%`,color=method)) +
  facet_grid(perf ~  n_rep,switch ="y") +
  geom_line() +
   geom_ribbon(color = NA,
    aes(x = threshold, ymin = `5%`, ymax = `95%`,fill=method),
     alpha = 0.5)  +
  scale_color_manual(values=alpha(res_colors,1)) +
    scale_x_continuous(breaks=c(0,0.25,0.5,0.75,1),
                     labels = c("0","25","50","75","100"))+
  scale_fill_manual(values=alpha(res_colors,0.6)) +
    scale_y_continuous(position = "right")+
   theme_bw() + theme(
                     legend.title = element_text(size=9),
                     legend.text = element_text(size=7,face = "italic"),
                     strip.background = element_rect(fill=alpha("gray",0.25)),
                  legend.background = element_rect(fill = alpha("white",0.1))) +
```

```
  guides(color=guide_legend(ncol=1,override.aes = list(size = 2))) +
  labs(y="",fill="Method",
      color="Method",x="Reproducibility threshold (%)")
p
```

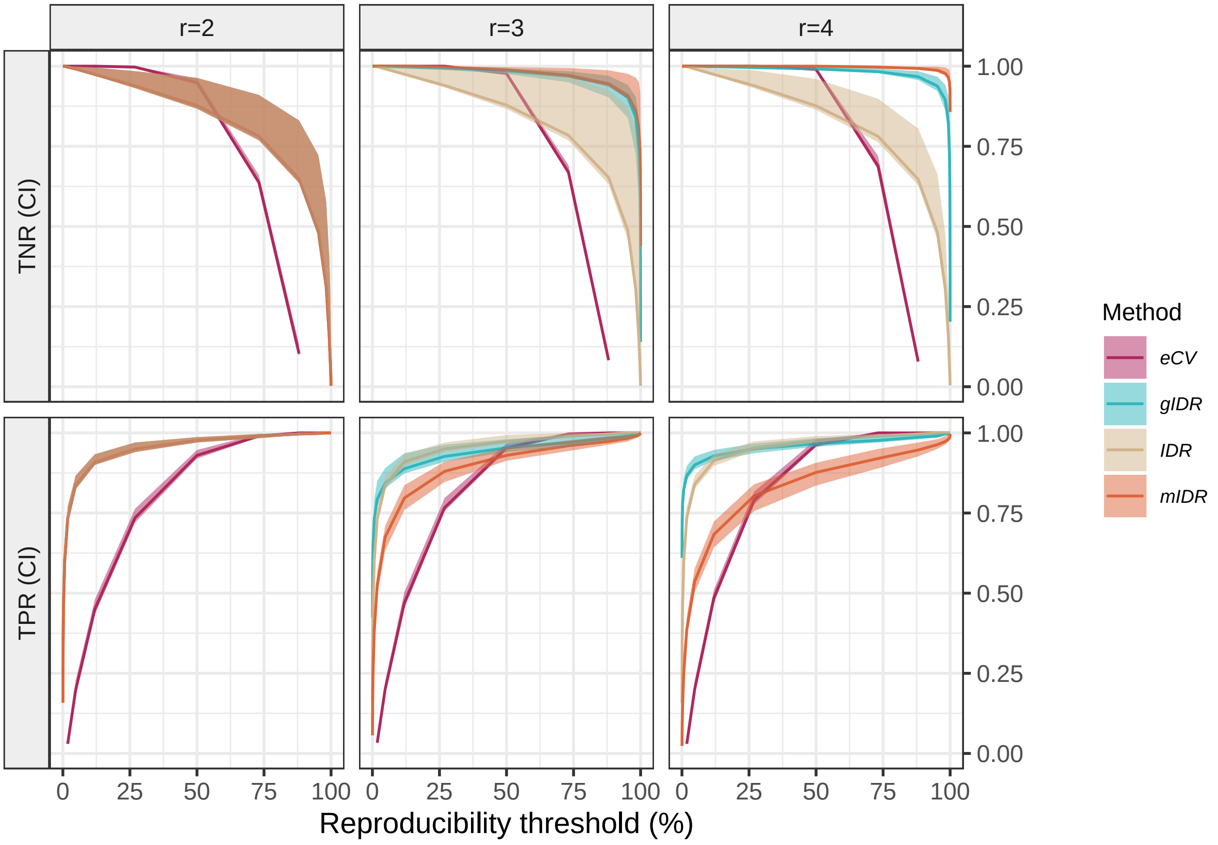


```
ggsave(filename = "Figure2.tiff",plot = p,device = "tiff",dpi=300,units = "in",
       width = 7,height = 5,scale = 0.85)
```

### Scenario 3.

```
p <- perf_plot_df %>%
  filter(scenario==3) %>%
  ggplot(aes(x=threshold,y=`50%`,color=method)) +
  facet_grid(perf ~  n_rep,switch ="y") +
  geom_line() +
   geom_ribbon(color = NA,
    aes(x = threshold, ymin = `5%`, ymax = `95%`,fill=method),
     alpha = 0.5)  +
  scale_color_manual(values=alpha(res_colors,1)) +
    scale_x_continuous(breaks=c(0,0.25,0.5,0.75,1),
                     labels = c("0","25","50","75","100"))+
  scale_fill_manual(values=alpha(res_colors,0.6)) +
    scale_y_continuous(position = "right")+
```

```
  theme_bw() + theme(
                  legend.title = element_text(size=9),
                  legend.text = element_text(size=7,face = "italic"),
                  strip.background = element_rect(fill=alpha("gray",0.25)),
               legend.background = element_rect(fill = alpha("white",0.1))) +
  guides(color=guide_legend(ncol=1,override.aes = list(size = 2))) +
  labs(y="",fill="Method",
      color="Method",x="Reproducibility threshold (%)")
p
```

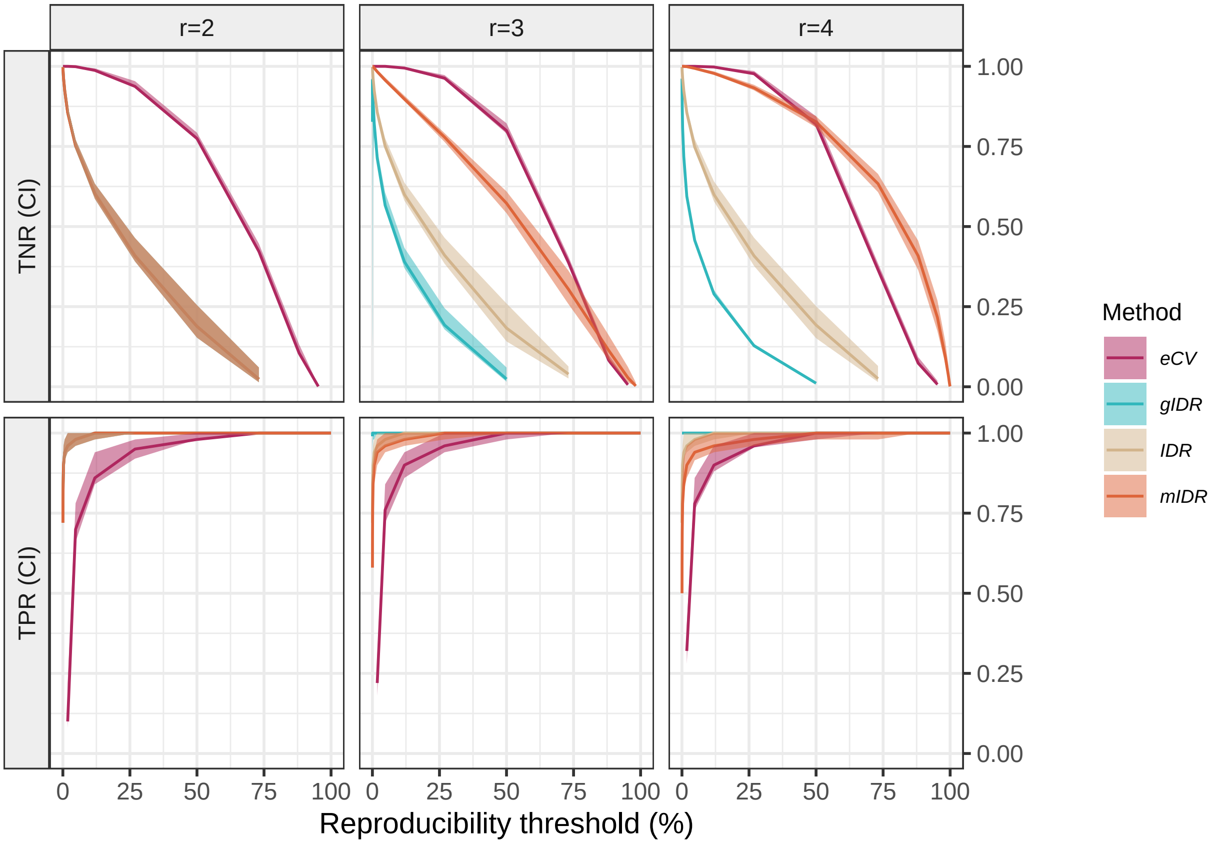


```
ggsave(filename = "Figure3.tiff",plot = p,device = "tiff",dpi=300,
       units = "in",width = 7,height = 5,scale = 0.85)
```
