## Supplementary File S2 for "eCV: Enhanced coefficient of variation and IDR extensions for reproducibility assessment of high-throughput experiments with multiple replicates"

### Experimental miR-eCLIP data from miR12424 and miR124 transfections

This document contains the required steps to generate the results for section *3.2.1. Experimental Scenario 1: miR-eCLIP data on transfected cells.*

Please follow the instructions in materials and methods within the main manuscript to download and process the experimental data. Pull a Docker image with all the required packages installed (`docker pull ecvpaper2024/ecv_results`).

### Data preparation.

Load required packages.

```
library("eCV")
Loading required package: idr
Loading required package: mvtnorm
Loading required package: future
Loading required package: future.apply
Loading required package: MatrixGenerics
Loading required package: matrixStats

Attaching package: 'MatrixGenerics'
The following objects are masked from 'package:matrixStats':

    colAlls, colAnyNAs, colAnys, colAvgsPerRowSet, colCollapse,
    colCounts, colCummaxs, colCummins, colCumprods, colCumsums,
    colDiffs, colIQRDiffs, colIQRs, colLogSumExps, colMadDiffs,
    colMads, colMaxs, colMeans2, colMedians, colMins, colOrderStats,
    colProds, colQuantiles, colRanges, colRanks, colSdDiffs, colSds,
    colSums2, colTabulates, colVarDiffs, colVars, colWeightedMads,
    colWeightedMeans, colWeightedMedians, colWeightedSds,
    colWeightedVars, rowAlls, rowAnyNAs, rowAnys, rowAvgsPerColSet,
    rowCollapse, rowCounts, rowCummaxs, rowCummins, rowCumprods,
    rowCumsums, rowDiffs, rowIQRDiffs, rowIQRs, rowLogSumExps,
    rowMadDiffs, rowMads, rowMaxs, rowMeans2, rowMedians, rowMins,
    rowOrderStats, rowProds, rowQuantiles, rowRanges, rowRanks,
    rowSdDiffs, rowSds, rowSums2, rowTabulates, rowVarDiffs, rowVars,
    rowWeightedMads, rowWeightedMeans, rowWeightedMedians,
    rowWeightedSds, rowWeightedVars



 --------------------------------------------------------------
|                                                          |
|                      ------    --                        |
|                ___  / ___\ \   / /                       |
|               / _ \ | |   \ \ / /                        |
```

```
|               |  __/ | |___  \ V /                         |
|                \___|  \____|  \_/                          |
|_________________________________________________________________|

        Enhanced Coefficient of Variation and IDR Extensions
                  for Reproducibility Assessment

This package provides extensions and alternative methods to  IDR  to
measure the reproducibility of omic data with an arbitrary number of
replicates. It introduces an enhanced Coefficient of Variation (eCV)
metric to assess the likelihood of omic features being reproducible.
library("idr2d")
require("tidyverse")
Loading required package: tidyverse
-- Attaching packages --------------------------------------- tidyverse 1.3.2 --
v ggplot2 3.4.3     v purrr   1.0.2
v tibble  3.2.1     v dplyr   1.1.2
v tidyr   1.3.0     v stringr 1.5.0
v readr   2.1.2     v forcats 0.5.1
-- Conflicts ------------------------------------------ tidyverse_conflicts() --
x dplyr::count()  masks matrixStats::count()
x dplyr::filter() masks stats::filter()
x dplyr::lag()    masks stats::lag()
library("future")
library("future.apply")
library("reshape2")

Attaching package: 'reshape2'

The following object is masked from 'package:tidyr':

    smiths
library("GenomicRanges")
Loading required package: stats4
Loading required package: BiocGenerics

Attaching package: 'BiocGenerics'

The following objects are masked from 'package:dplyr':

    combine, intersect, setdiff, union

The following objects are masked from 'package:stats':

    IQR, mad, sd, var, xtabs

The following objects are masked from 'package:base':

    anyDuplicated, append, as.data.frame, basename, cbind, colnames,
    dirname, do.call, duplicated, eval, evalq, Filter, Find, get, grep,
    grepl, intersect, is.unsorted, lapply, Map, mapply, match, mget,
    order, paste, pmax, pmax.int, pmin, pmin.int, Position, rank,
    rbind, Reduce, rownames, sapply, setdiff, sort, table, tapply,
```

```
    union, unique, unsplit, which.max, which.min

Loading required package: S4Vectors

Attaching package: 'S4Vectors'

The following objects are masked from 'package:dplyr':

    first, rename

The following object is masked from 'package:tidyr':

    expand

The following objects are masked from 'package:base':

    expand.grid, I, unname

Loading required package: IRanges

Attaching package: 'IRanges'

The following objects are masked from 'package:dplyr':

    collapse, desc, slice

The following object is masked from 'package:purrr':

    reduce

Loading required package: GenomeInfoDb
library("BSgenome")
Loading required package: Biostrings
Loading required package: XVector

Attaching package: 'XVector'

The following object is masked from 'package:purrr':

    compact


Attaching package: 'Biostrings'

The following object is masked from 'package:base':

    strsplit

Loading required package: rtracklayer
library("BSgenome.Hsapiens.UCSC.hg38")
library("Biostrings")
library('pROC')
Type 'citation("pROC")' for a citation.
```

```
Attaching package: 'pROC'

The following objects are masked from 'package:IRanges':

    cov, var

The following objects are masked from 'package:S4Vectors':

    cov, var

The following object is masked from 'package:BiocGenerics':

    var

The following objects are masked from 'package:stats':

    cov, smooth, var
library('microRNA')
library("DESeq2")
Loading required package: SummarizedExperiment
Loading required package: Biobase
Welcome to Bioconductor

    Vignettes contain introductory material; view with
    'browseVignettes()'. To cite Bioconductor, see
    'citation("Biobase")', and for packages 'citation("pkgname")'.


Attaching package: 'Biobase'

The following object is masked from 'package:MatrixGenerics':

    rowMedians

The following objects are masked from 'package:matrixStats':

    anyMissing, rowMedians
library("DescTools")
```

Set color palette.

```
res_colors <-
  c(IDR = "tan",
    gIDR = "#30B7BC", # bright teal
    eCV = "#AF275F", # light magenta
    mIDR = "#DE653A" # medium teal
  )
```

Upload miR-eCLIP data. Use only chromosome 1.

```
(mireclip_data <-
  read_tsv(
```

```
    file = "miR1_miR124_transfections.miR_chim_peak_table.tsv",
    col_types = cols()) %>%
   dplyr::filter(chrom == "chr1")
  )
# A tibble: 4,059 x 12
   chrom  start    end strand gene     ensg               feature    Symbol_miRNA
   <chr>  <dbl>  <dbl> <chr>  <chr>    <chr>              <chr>      <chr>
 1 chr1  632106 632157 +      MTCO1P12 ENSG00000237973.1 noncoding~ hsa-miR-1-3p
 2 chr1  632106 632157 +      MTCO1P12 ENSG00000237973.1 noncoding~ hsa-miR-124~
 3 chr1  632106 632205 +      MTCO1P12 ENSG00000237973.1 noncoding~ hsa-miR-1-3p
 4 chr1  632106 632205 +      MTCO1P12 ENSG00000237973.1 noncoding~ hsa-miR-124~
 5 chr1  633955 634060 +      MTATP6P1 ENSG00000248527.1 noncoding~ hsa-miR-1-3p
 6 chr1  633955 634060 +      MTATP6P1 ENSG00000248527.1 noncoding~ hsa-miR-124~
 7 chr1  633955 634060 +      MTATP6P1 ENSG00000248527.1 noncoding~ hsa-miR-182~
 8 chr1  633955 634060 +      MTATP6P1 ENSG00000248527.1 noncoding~ hsa-miR-196~
 9 chr1  633955 634060 +      MTATP6P1 ENSG00000248527.1 noncoding~ hsa-miR-20a~
10 chr1  633955 634060 +      MTATP6P1 ENSG00000248527.1 noncoding~ hsa-miR-378~
# i 4,049 more rows
# i 4 more variables: At13_CaptureT1_miR1_rep1 <dbl>,
#   At14_CaptureT1_miR1_rep2 <dbl>, At15_CaptureT124_miR124_rep1 <dbl>,
#   At16_CaptureT124_miR124_rep2 <dbl>
```

Extract peaks intensities.

```
mireclip_inten <-
  mireclip_data %>%
  dplyr::select(contains("Capture"))
```

Filter peaks with zero intensity in more than one replicate.

```
tmp <- rowSums(mireclip_inten >=
                   3) > 0
mireclip_data <- mireclip_data[tmp,]
mireclip_inten <- mireclip_inten[tmp,]
```

```
mireclip_inten <-
  idr2d::preprocess(x = mireclip_inten %>%
              mutate_all(~ . + 1.1) %>%
               as.matrix(),
             value_transformation = "identity",
            jitter_factor = 0) %>%
  log()
```

Rename features.

```
# Rename features.
region_rename <- c(
  "5utr" = "5' UTR",
  "CDS" = "CDS",
  "3utr" = "3' UTR",
  "noncoding_exon" = "Other",
  "miRNA" = "miRNA",
```

```
  "miRNA_proximal" = "miRNA",
  "5ss" = "Intron",
  "noncoding_5ss" = "Other",
  "3ss" = "Intron",
  "noncoding_3ss" = "Other",
  "proxintron" = "Intron",
  "noncoding_proxintron" = "Other",
  "distintron" = "Intron",
  "noncoding_distintron" = "Other",
  "tRNA" = "Other",
  "intergenic" = "Other"
)

mireclip_data$feature <-
  factor(x = mireclip_data$feature,
         levels = names(region_rename),
         labels = region_rename)
```

```
# Subset conditions.
conditions <- grep(colnames(mireclip_inten), pattern = "_miR124_")
mireclip_inten_reps <-
  list( "miR1" = mireclip_inten[, -conditions],
       "miR124" = mireclip_inten[, conditions])

# Check dimensions.
lapply(mireclip_inten_reps, dim)
$miR1
[1] 1858    2

$miR124
[1] 1858    2
```

Extract mean intensities by peaks across replicates.

```
mireclip_inten_means <- rowMeans(mireclip_inten, na.rm = TRUE)
```

Set initial values for each method.

```
# Set parameters for each model.
methods_params <- list(
  eCV = list(max.ite = 1e4),
  gIDR = list(
     mu = 2,
     sigma = 5,
     rho = 0.95,
     p = 0.7,
    eps = 1e-3,
    max.ite = 50
  ),
  IDR = list(
     mu = 2,
     sigma = 5,
     rho = 0.95,
```

```
    p = 0.7,
    eps = 1e-3,
    max.ite = 50
  ),
  mIDR = list(
    mu = 2,
    sigma = 5,
    rho = 0.95,
    p = 0.7,
    eps = 1e-3,
    max.ite = 50
  )
)
```

### Data analysis.

#### Intensity of chimeric peaks involving transfected miRNAs versus endogenous chimeras.

```
boxplot(c(rowMeans(mireclip_inten_reps$miR1),rowMeans(mireclip_inten_reps$miR124)) ~
          c(mireclip_data$Symbol_miRNA == "hsa-miR-1-3p",
          mireclip_data$Symbol_miRNA == "hsa-miR-124-3p"), xlab="Peaks with transfected miRNAs",ylab="Pe
```

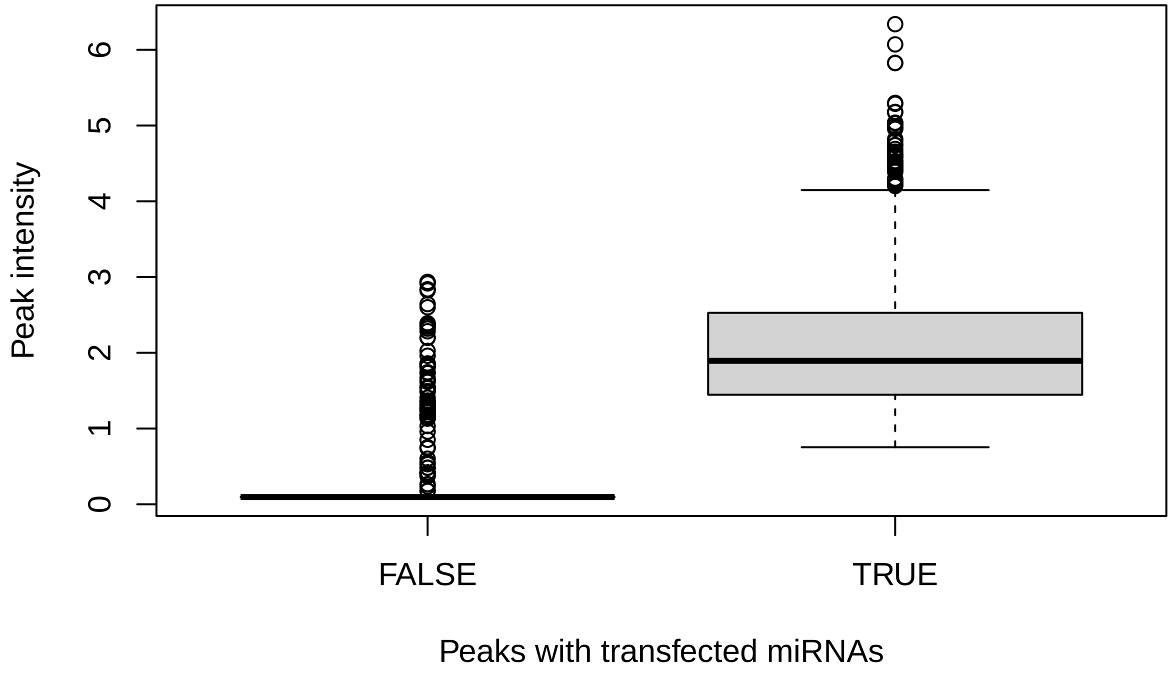

#### Correlation analysis of impossed reproducible features.

```
R1 <- cor(mireclip_inten[mireclip_data$Symbol_miRNA == "hsa-miR-124-3p", 3:4])
R2 <- cor(mireclip_inten[mireclip_data$Symbol_miRNA == "hsa-miR-1-3p", 1:2],
          use = "pairwise.complete.obs")

r1 <- R1[upper.tri(R1)]
r2 <- R2[upper.tri(R2)]

CorCI(r1,sum(mireclip_data$Symbol_miRNA == "hsa-miR-124-3p"))
      cor    lwr.ci    upr.ci
0.8149195 0.7975457 0.8309426
CorCI(r2,sum(mireclip_data$Symbol_miRNA == "hsa-miR-1-3p"))
      cor    lwr.ci    upr.ci
0.6287137 0.5485516 0.6974230
```

#### Assess reproducibility with each method.

Create a table with all combinations of parameters.

```
par_settings <- expand.grid(n_rep=c("miR1","miR124"),
                            method=c("IDR","eCV"),
                            stringsAsFactors = FALSE) %>%
  mutate(par_com = seq_along(method))
```

```
# Set parallel scheduler.
future::plan(multisession, workers = 4)
perf_res <- NULL
for (i in par_settings$par_com) {
  set.seed(42)
  print(par_settings[i,])
  method <- par_settings$method[i]
  n_rep <- par_settings$n_rep[i]
  if(method != "eCV") {
    X <- preprocess(mireclip_inten_reps[[n_rep]],
                        value_transformation = "identity",
                        jitter=1e-5)
  } else {
    X <- mireclip_inten_reps[[n_rep]]
  }

  rep_index <- mrep_assessment(
          x = X,
          method = method,
          param = methods_params[[method]],
          n_threads = 4
        )$rep_index

   tmp1 <- mireclip_data$Symbol_miRNA == "hsa-miR-124-3p"
   tmp2 <- mireclip_data$Symbol_miRNA == "hsa-miR-1-3p"

   perf <- roc(tmp1 | tmp2, rep_index, quiet = TRUE)
```

```
   perf_thr <- ci.coords(perf,conf.level=0.90,
                         x =  1/(1 + exp(-c(1:20) + 10)),
                         ret=c("threshold","tpr","tnr"))
    perf_thr <- rbind(perf_thr$tpr %>%
    as.data.frame() %>%
    mutate(threshold= 1/(1 + exp(-c(1:20) + 10)),
           perf="TPR (CI)") ,
        perf_thr$tnr %>%
    as.data.frame() %>%
    mutate(threshold= 1/(1 + exp(-c(1:20) + 10)),
           perf="TNR (CI)"))

  perf_thr$method <- method
  perf_thr$n_rep <- n_rep
  perf_res <- rbind(perf_res, perf_thr)
}

# Close me buddies.
future::plan(sequential)

# Save results.
saveRDS(perf_res,file="perf_resRealTransf.rds")
```

Arrange results for figure creation.

```
p <- perf_res %>%
  mutate(n_rep=factor(n_rep,levels = c("miR124","miR1"), ordered = TRUE)) %>%
  ggplot(aes(x=threshold,y=`50%`,color=method)) +
  facet_grid(perf~n_rep,switch ="y") +
   geom_ribbon(color = NA,
    aes(x = threshold, ymin = `5%`, ymax = `95%`,fill=method),
     alpha = 0.3)  +
   geom_line() +
  scale_color_manual(values=alpha(res_colors,1)) +
  scale_fill_manual(values=alpha(res_colors,0.6)) +
    scale_y_continuous(position = "right")+
   theme_bw() + theme(
                     legend.title = element_text(size=9),
                     legend.text = element_text(size=7,face = "italic"),
                     strip.background = element_rect(fill=alpha("gray",0.25)),
                  legend.background = element_rect(fill = alpha("white",0.1))) +
   guides(color=guide_legend(ncol=1,override.aes = list(size = 2))) +
   labs(y="",fill="Method",linetype="Transfection",
       color="Method",x="Reproducibility threshold (%)") +
   scale_x_continuous(breaks=c(0,0.25,0.5,0.75,1),
                     labels = c("0","25","50","75","100"))

p
```

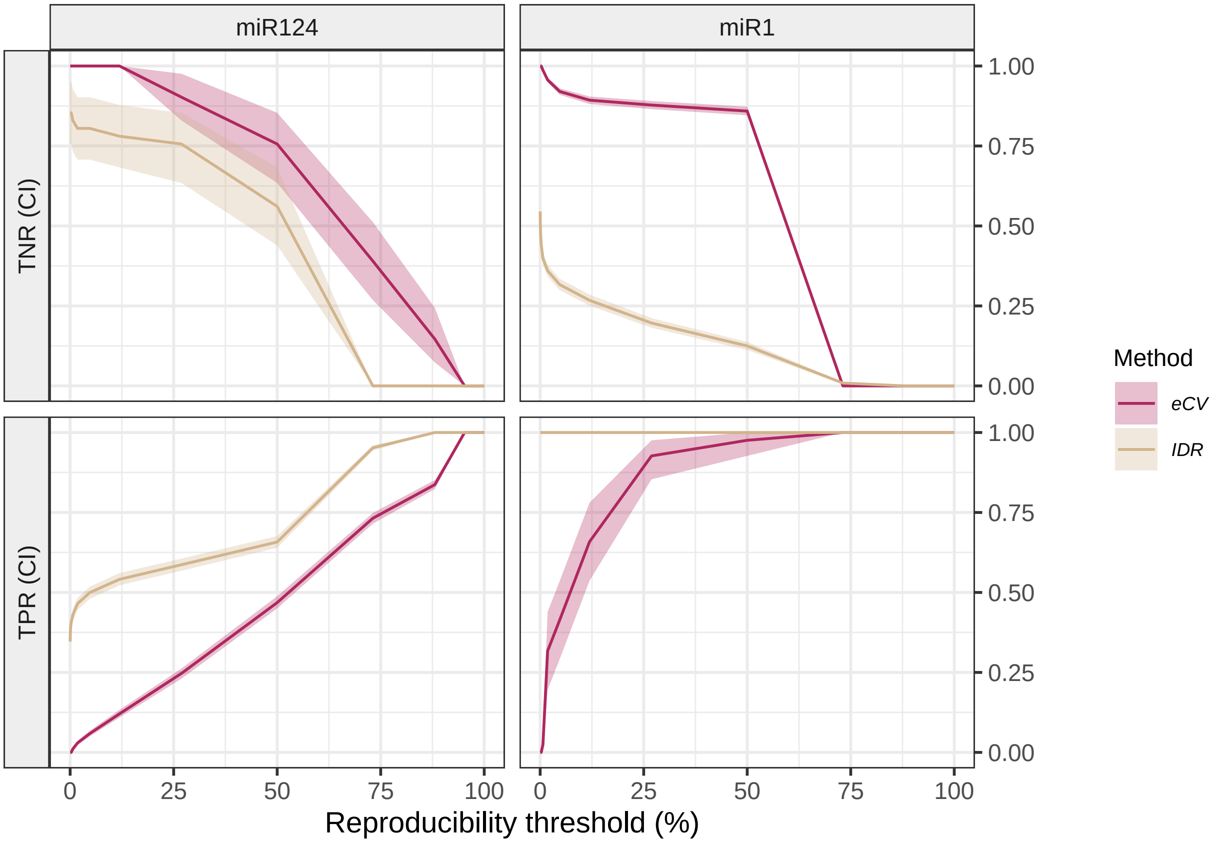


```
ggsave(filename = "Figure4.tiff",plot = p,device = "tiff",
       dpi=300,units = "in",width = 7,height = 5,scale = 0.85)
```
