## Supplementary File S3 for "eCV: Enhanced coefficient of variation and IDR extensions for reproducibility assessment of high-throughput experiments with multiple replicates"

### Experimental miR-eCLIP data from wild-type cells

This document contains the required steps to generate the results for section *3.2.2: Experimental Scenario 2: miR-eCLIP data from wild-type cells.*

Set color palette.

```
res_colors <-
  c(IDR = "tan",
    gIDR = "#30B7BC",
    eCV = "#AF275F", # light magenta
    mIDR = "#DE653A" # medium teal
  )
```

Upload miR-eCLIP data and keep only chromosome 1.

```
(mireclip_data <-
  read_tsv(
    file = "CH010_merged_peaks.miR_chim_peak_table.tsv",
    col_types = cols()) %>%
   dplyr::filter(chrom == "chr1")
```

```
  )
# A tibble: 137,259 x 16
   chrom start   end strand gene                          ensg  feature Symbol_miRNA
   <chr> <dbl> <dbl> <chr>  <chr>                         <chr> <chr>   <chr>
 1 chr1  89749 89840 -      ENSG00000238009|ENSG0000~ ENSG~ noncod~ hsa-miR-17-~
 2 chr1  89749 89840 -      ENSG00000238009|ENSG0000~ ENSG~ noncod~ hsa-miR-193~
 3 chr1  89749 89840 -      ENSG00000238009|ENSG0000~ ENSG~ noncod~ hsa-miR-20a~
 4 chr1  89749 89840 -      ENSG00000238009|ENSG0000~ ENSG~ noncod~ hsa-miR-223~
 5 chr1  89749 89840 -      ENSG00000238009|ENSG0000~ ENSG~ noncod~ hsa-miR-25-~
 6 chr1  89749 89840 -      ENSG00000238009|ENSG0000~ ENSG~ noncod~ hsa-miR-27a~
 7 chr1  89749 89840 -      ENSG00000238009|ENSG0000~ ENSG~ noncod~ hsa-miR-92a~
 8 chr1  89749 89840 -      ENSG00000238009|ENSG0000~ ENSG~ noncod~ hsa-miR-92b~
 9 chr1  89840 89922 -      ENSG00000239945|ENSG0000~ ENSG~ noncod~ hsa-miR-17-~
10 chr1  89840 89922 -      ENSG00000239945|ENSG0000~ ENSG~ noncod~ hsa-miR-20a~
# i 137,249 more rows
# i 8 more variables: CH010_1_IP1_S17_L001_R1_001 <dbl>,
#   CH010_2_IP2_S18_L001_R1_001 <dbl>, CH010_3_IP3_S19_L001_R1_001 <dbl>,
#   CH010_4_IP4_S20_L001_R1_001 <dbl>, CH010_5_IP5_S21_L001_R1_001 <dbl>,
#   CH010_6_IP6_S22_L001_R1_001 <dbl>, CH010_7_IP7_S23_L001_R1_001 <dbl>,
#   CH010_8_IP8_S24_L001_R1_001 <dbl>
```

### Data analysis.

Filter out peaks without enough counts.

```
tmp <-
  (mireclip_data %>%
  dplyr::select(contains("CH010")) %>% rowMeans()) > 3
tmp <-
mireclip_data <- mireclip_data[tmp, ]
```

Get PCA of peak intensities.

```
pca <-
  mireclip_data %>%
  dplyr::select(contains("CH010")) %>%
  ceiling() %>% mutate_all(~ log(. + 1)) %>%
  princomp()
ggplot(pca$loadings[,TRUE] %>% as.data.frame() %>% rownames_to_column("Sample"),
       aes(x=Comp.1, y=Comp.2)) +
  ggrepel::geom_label_repel(aes(label=Sample),size=2) + geom_point() +
  theme_bw() + ggtitle("PCA biplot of peaks intensities")
```

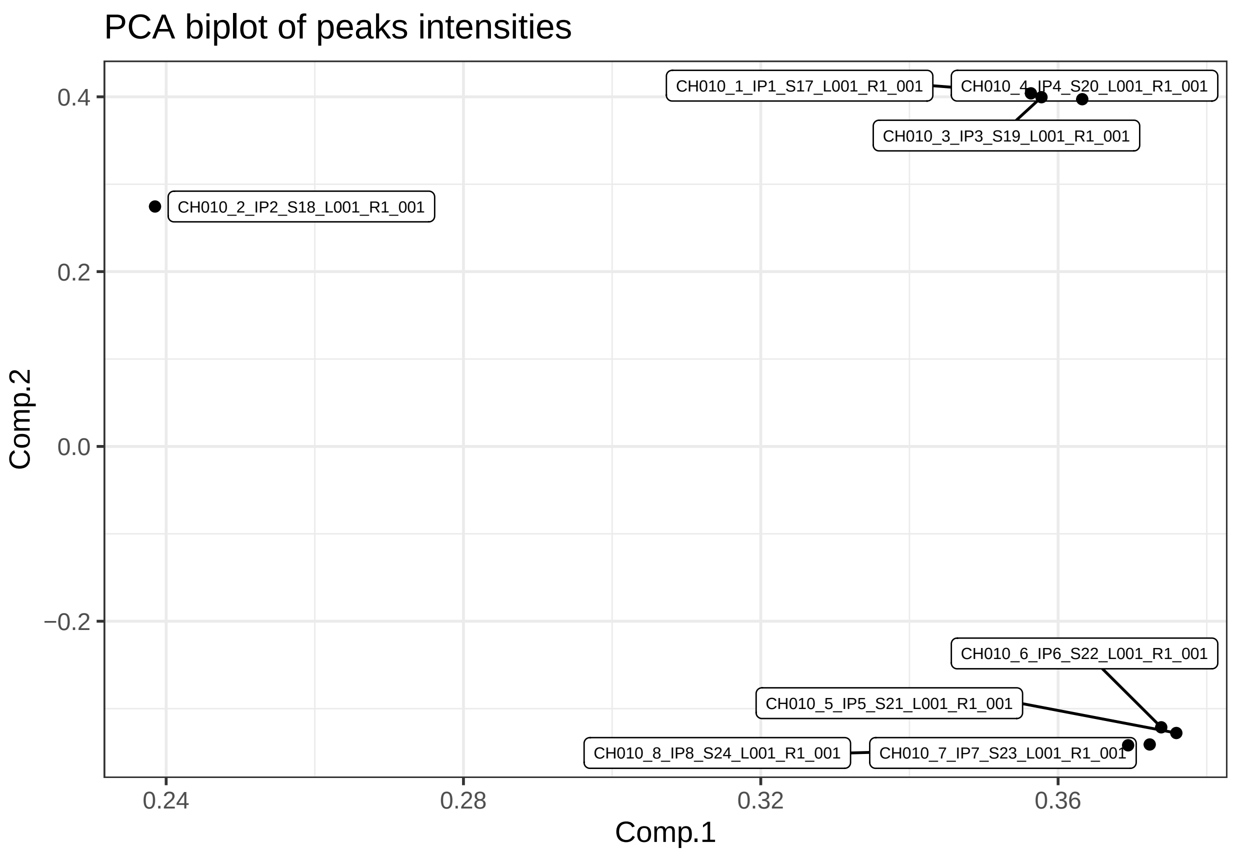


Create subset with different number of replicates and use four replicates within the same condition.

```
# Subset replicates.
mireclip_data <- mireclip_data %>%
  dplyr::select(-rownames(pca$loadings[pca$loadings[, 2] > 0, ]))
```

Use PCs to obtain number of independent groups of peaks for multiple comparisons adjustment.

```
pc <- pca$scores

# Function to convert binary vector to decimal number
binary_to_decimal <- function(binary_vector) {
  decimal_number <- sum(binary_vector * 2^(rev(seq_along(binary_vector) - 1)))
      return(decimal_number)
}
# We take scores at, or above, the third Quartile as "informative".
# The remaining scores are taken as "uninformative".
# Number of clusters is estimated by turning binary rows into
# single decimals numbers.
decimal_numbers <- apply(
      apply(pc, 2, function(x) as.numeric(x >= quantile(x, probs = 0.75))),
      1, binary_to_decimal
)

# The estimated number of independent tests is used to get.
# The number of independent tests is estimated by taking the number of
```

```r
    # different decimal numbers.
n_ind_peaks <- sum(unique(decimal_numbers) != 0)
```

#### Seed matching analysis.

Create a genomic range object to extract sequence information.

```r
# Create a variable with formatted information
mireclip_data <- mireclip_data %>%
  mutate(genomic_info = paste0(chrom,
                               ":",
  start,
  "-",
  end,
  strand),)
gr <- makeGRangesFromDataFrame(mireclip_data,
                               keep.extra.columns = TRUE)
unique_peaks <- unique(gr)
```

Extract peaks sequences.

```r
(peak_seqs <- getSeq(BSgenome.Hsapiens.UCSC.hg38, unique_peaks))
DNAStringSet object of length 1897:
       width seq
   [1]    91 ATACCACCAATCAATACTCATCATTAATAATC...ATAGCCCCCTTTCACTTCTGAGTCCCAGAGGT
   [2]    84 CTGTAGGCTCATTCATTTCTCTAACAGCAGTA...TTGAGAAGCCTTCGCTTCGAAGCGAAAAGTCC
   [3]   100 AACCACCCAACTATCTATAAACCTAGCCATGG...TTCGCTCTAAGATTAAAAATGCCCTAGCCCAC
   [4]    72 AATTCTACTGACTATCCTAGAAATCGCTGTCG...CAAGCCTACGTTTTTACACTTCTAGTAAGCCT
   [5]   117 CCTGAACTTCCCTGAGATCAAACGAAGGAAGA...AACAGCTCTGAAGAGGACGACACCGAGGGATT
   ...   ... ...
[1893]    29 GACAAGACATCACAGTGGTCTGGGCTGGA
[1894]   114 AGGAACTGAACCCCTCAGGATCCCCGCCGACC...ATTCTCCAGCTCACTCCCAATCCCAGGCTCAC
[1895]    91 GTTGCCAATTGTATCTGTTTTTATGAAATGTT...ATGGGGAGACCATAGAAGAATGATCCAAGGAG
[1896]   106 AAGGATGTTCTGGGAAACCTCTCCCGATTCAC...AACTAATCTTCTCATACTTACATTTTGCAGAT
[1897]   115 ATGCTTCAGGGAGTACCACATCCGGTGACATC...TGGCAGTTGAGCACTTCTGTTTTGTGTTGGAA
names(peak_seqs) <- unique_peaks$genomic_info
```

Extract peaks intensities.

```r
mireclip_inten <-
  mireclip_data %>%
  dplyr::select(contains("CH010"))
```

```r
N_counts <-
  mireclip_data %>% dplyr::select(contains("CH010"),
                                  MIRNA=Symbol_miRNA,ENSG=ensg)
peak_cols <- grep(colnames(N_counts),
                  pattern="CH010")

N_counts[,peak_cols] <- ceiling(N_counts[,peak_cols]) + 1
```

```
# Get total chimeric counts.
n <- N_counts[, peak_cols] %>% colSums() %>%
    as.vector() %>%
    as.numeric()

ind_test0 <- function(x,dist="nbinom") {

  f <-
    list('binom'=function(nij,pi,pj,n) pbinom(q = nij,
                                              size=n,
                                              prob = pi*pj,
                                              lower.tail = T),
         'nbinom'=function(nij,pi,pj,n) {
             pnbinom(q = nij,size= 1,
                     prob = pi*pj,
                                              lower.tail = T)

         }
  )


  res <- future_lapply(seq_len(nrow(x)), function(k) {
    gene_i <- x$ENSG[k]
    mirna_j <- x$MIRNA[k]

  # Get chimeric counts.
  nij <- x[k, peak_cols] %>%
    as.vector() %>%
    as.numeric()

  # Get marginal gene and mirna counts.
  ni <- x[x$ENSG == gene_i ,
                      peak_cols] %>%
    colSums() %>%
    as.vector() %>%
    as.numeric()

  nj <- x[x$MIRNA == mirna_j,
                      peak_cols] %>%
    colSums()%>%
    as.vector() %>%
    as.numeric()

    # Get marginal probabilities.
    pij <- nij / n
    pi <- ni / n
    pj <- nj / n

  pvalue <-
   f[[dist]](nij,pi,pj,n)

  return(pvalue)
}) %>% do.call(what = rbind)
```

```
  return(res)
}
future::plan(multisession, workers = 4)
chim_test <- ind_test0(N_counts)
# Close me buddies.
future::plan(sequential)

mireclip_inten <-
  idr2d::preprocess(x = mireclip_inten %>%
             mutate_all(~ . + 1.1) %>%
              as.matrix(),
            value_transformation = "identity",
           jitter_factor = 0) %>%
  log()
```

Represent peaks based on most enriched miRNA.

```
sig_enriched_mirnas <-
  lapply(unique_peaks$genomic_info, function (peak) {
  cat(".")
  tmp <- which(mireclip_data$genomic_info == peak)
  if (length(tmp) == 1) {
    mu <- mean(mireclip_inten[tmp,])
    sd <- sd(mireclip_inten[tmp,])
    mirna_pval <- pt(2 * mu/sd, lower.tail = FALSE, df = 3)
    mirna_padj <- mirna_pval * n_ind_peaks
    mirna_padj <-
      ifelse(mirna_padj>= 1, 0.99, mirna_padj)

    if (mirna_padj < 0.1)
        return(mireclip_data$Symbol_miRNA[tmp])
   else  NA
  }
  else {
    mu <- rowMeans(mireclip_inten[tmp,])
    sd <- MatrixGenerics::rowSds(mireclip_inten[tmp,])
    mirna_pval <- pt(2 * mu/sd, lower.tail = FALSE, df = 3)
    mirna_padj <- mirna_pval * n_ind_peaks
    mirna_padj <-
      ifelse(mirna_padj>= 1, 0.99, mirna_padj)

    if (any(mirna_padj < 0.1))
        return(mireclip_data$Symbol_miRNA[tmp][mirna_padj < 0.05])
    else NA
  }
})

with_sig_mirna <-
  !(lapply(sig_enriched_mirnas, is.na) %>%
      unlist())
names(with_sig_mirna) <-
  names(sig_enriched_mirnas) <-
  unique_peaks$genomic_info
```

```
mireclip_data$with_sig_mirna <- with_sig_mirna[mireclip_data$genomic_info]
```

Rename features.

```
# Rename features.
region_rename <- c(
  "5utr" = "5' UTR",
  "CDS" = "CDS",
  "3utr" = "3' UTR",
  "noncoding_exon" = "Other",
  "miRNA" = "miRNA",
  "miRNA_proximal" = "miRNA",
  "5ss" = "Intron",
  "noncoding_5ss" = "Other",
  "3ss" = "Intron",
  "noncoding_3ss" = "Other",
  "proxintron" = "Intron",
  "noncoding_proxintron" = "Other",
  "distintron" = "Intron",
  "noncoding_distintron" = "Other",
  "tRNA" = "Other",
  "intergenic" = "Other"
)

mireclip_data$feature <-
  factor(x = mireclip_data$feature,
         levels = names(region_rename),
         labels = region_rename)
```

Get peaks with positive seed matching using scanMiR.

```
# Get peak strand.
strand <- as.character(strand(unique_peaks))
names(strand) <- unique_peaks$genomic_info

# Get affinity constants.
KSmodels <- getKdModels(species = "hsa")

seed_match_res <-
lapply(names(with_sig_mirna[with_sig_mirna]),
                function(peak) {
                  cat(".")
    # Get miRNA from enriched peaks.
    mirnas <- sig_enriched_mirnas[[peak]]

    peak_seqs_chr <- as.character(ifelse(strand[peak] == "+",
                            peak_seqs[peak],
                            reverseComplement(peak_seqs[peak])))

    # Does the peak sequence matches seed?
    seed_match <-
      try(aggregateMatches(findSeedMatches(seqs = peak_seqs_chr,
                seeds = KSmodels[mirnas],
```

```
                  p3.extra = TRUE,
                  onlyCanonical = FALSE,
                  minDist = -Inf,
                  verbose = FALSE,maxLogKd = Inf)), silent = TRUE)
  return(seed_match)
})

names(seed_match_res) <-
  names(with_sig_mirna[with_sig_mirna])

with_seed_match <- (lapply(seed_match_res, class) %>% unlist()) != "try-error"

mireclip_data$with_seed_match <- FALSE
mireclip_data$with_seed_match[mireclip_data$genomic_info %in%
                              names(with_seed_match)[with_seed_match]] <-
  with_seed_match
```

Create subset with different number of replicates.

```
# Subset replicates.
mireclip_inten_reps <-
  list("n_rep=2" = mireclip_inten[, 1:2],
       "n_rep=3" = mireclip_inten[, 1:3],
       "n_rep=4" = mireclip_inten[, 1:4])

# Check dimensions.
lapply(mireclip_inten_reps, dim)
$`n_rep=2`
[1] 3145    2

$`n_rep=3`
[1] 3145    3

$`n_rep=4`
[1] 3145    4
```

Set initial values for each method.

```
# Set parameters for each model.
methods_params <- list(
  eCV = list(max.ite = 1e4),
  gIDR = list(
     mu = 2,
     sigma = 1,
     rho = 0.5,
     p = 0.5,
    eps = 1e-3,
    max.ite = 100
  ),
  IDR = list(
     mu = 2,
     sigma = 1,
     rho = 0.5,
```

```
    p = 0.5,
    eps = 1e-3,
    max.ite = 100
  ),
  mIDR = list(
    mu = 2,
    sigma = 1,
    rho = 0.5,
    p = 0.5,
    eps = 1e-3,
    max.ite = 100
  )
)
```

#### Correlation analysis of impossed reproducible features.

```
tmp <-
     mireclip_data$with_seed_match

mireclip_data$Symbol_miRNA[tmp] %>% unique() %>% length()
r <- lapply(unique(mireclip_data$genomic_info[tmp]),function(x) {
  i <- which(mireclip_data$genomic_info[tmp] == x)
  if (length(i) > 1) {
    R<- cor(mireclip_inten[tmp, ][i,])
    r <- mean(R[upper.tri(R)])
    m <- mean(mireclip_inten[tmp, ][i,])
c(length(i),r,m)
      } else {c(NA,NA,NA)}})

d <- do.call(rbind,r) %>% as.data.frame() %>% drop_na()
```

#### Assess reproducibility with each method.

Create a table with all combinations of parameters.

```
(par_settings <- expand.grid(n_rep =  paste0("n_rep=",4:2),
                            method=c("gIDR","mIDR","IDR","eCV"),
                            stringsAsFactors = FALSE) %>%
  mutate(par_com = seq_along(n_rep)))
     n_rep method par_com
1  n_rep=4   gIDR       1
2  n_rep=3   gIDR       2
3  n_rep=2   gIDR       3
4  n_rep=4   mIDR       4
5  n_rep=3   mIDR       5
6  n_rep=2   mIDR       6
7  n_rep=4    IDR       7
8  n_rep=3    IDR       8
9  n_rep=2    IDR       9
10 n_rep=4    eCV      10
```

```
11 n_rep=3    eCV      11
12 n_rep=2    eCV      12
```

```
# Set parallel scheduler.
future::plan(multisession, workers = 4)
perf_res <- NULL
for (i in par_settings$par_com) {
  set.seed(42)
  print(par_settings[i,])
  n_rep <- par_settings$n_rep[i]
  method <- par_settings$method[i]

  if(method != "eCV") {
    X <- preprocess(mireclip_inten_reps[[n_rep]],
                         value_transformation = "identity",
                         jitter = 1e-4)
  } else {
    X <- mireclip_inten_reps[[n_rep]]
  }

  rep_index <- mrep_assessment(
          x = X,
          method = method,
          param = methods_params[[method]],
          n_threads = 4
        )$rep_index

   tmp <- exp(rowMeans(log(chim_test))) > 0.01 &
     mireclip_data$with_seed_match | mireclip_data$feature == "3'UTR"

   print(perf <- roc(tmp,
              rep_index, quiet = TRUE))

  perf_thr <- ci.coords(perf,conf.level=0.90,
                        x =  1/(1 + exp(-c(1:20) + 10)),
                        ret=c("threshold","tpr","tnr"))
    perf_thr <- rbind(perf_thr$tpr %>%
    as.data.frame() %>%
    mutate(threshold= 1/(1 + exp(-c(1:20) + 10)),
           perf="TPR (CI)") ,
        perf_thr$tnr %>%
    as.data.frame() %>%
    mutate(threshold= 1/(1 + exp(-c(1:20) + 10)),
           perf="TNR (CI)"))

  perf_thr$n_rep <- n_rep
  perf_thr$method <- method
  perf_res <- rbind(perf_res, perf_thr)
}

# Close me buddies.
future::plan(sequential)
```

```r
# Save results.
saveRDS(perf_res,file="perf_resRealmiReCLIP.rds")
```

Arrange results for figure creation.

```r
p <- perf_res %>%
  mutate(n_rep = paste0("r=",str_remove(n_rep,"n_rep="))) %>%
  ggplot(aes(x=threshold,y=`50%`,color=method)) +
  facet_grid(perf~n_rep,switch ="y") +
   geom_ribbon(color = NA,
    aes(x = threshold, ymin = `5%`, ymax = `95%`,fill=method),
     alpha = 0.3)  +
   geom_line() +
  scale_color_manual(values=alpha(res_colors, 1)) +
  scale_linetype_manual(values=c("miR1"="solid", "miR124"="dashed")) +
  scale_fill_manual(values=alpha(res_colors, 0.6)) +
    scale_y_continuous(position = "right")+
   theme_bw() + theme(
                     legend.title = element_text(size=9),
                     legend.text = element_text(size=7, face = "italic"),
                     strip.background = element_rect(fill=alpha("gray", 0.25)),
                  legend.background = element_rect(fill = alpha("white", 0.1))) +
   guides(color=guide_legend(ncol=1,override.aes = list(size = 2))) +
   labs(y="",fill="Method",linetype="Transfection",
       color="Method",x="Reproducibility threshold (%)") +
   scale_x_continuous(breaks=c(0, 0.25, 0.5, 0.75, 1),
                     labels = c("0","25","50","75","100"))

p
```

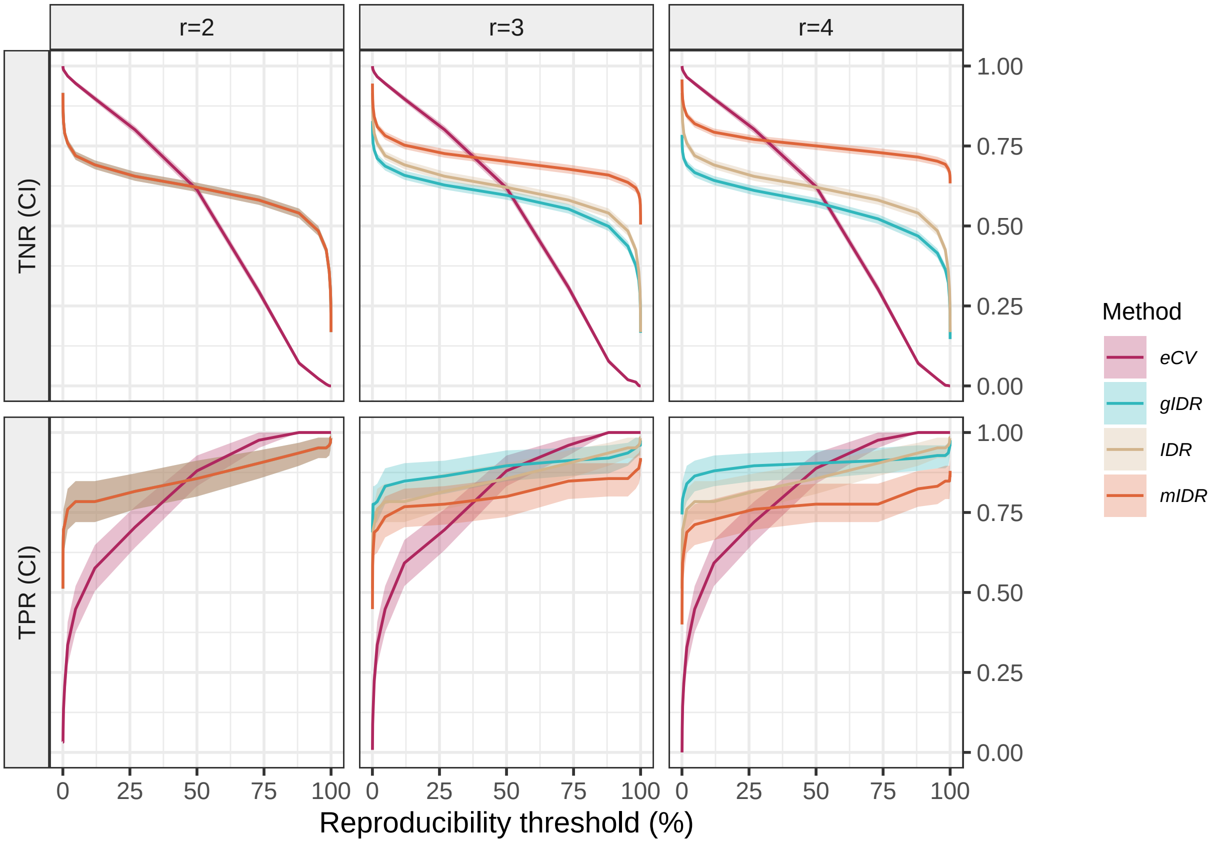


```
ggsave(filename = "Figure5.tiff",plot = p,device = "tiff",
       dpi=300,units = "in",width = 7,height = 5,scale = 0.85)
```
