## Supplementary File S4 for "eCV: Enhanced coefficient of variation and IDR extensions for reproducibility assessment of high-throughput experiments with multiple replicates"

### Experimental eCLIP data from wild-type cells

This document contains the required steps to generate the results for section *3.2.3: Experimental Scenario 3: RBFOX2 eCLIP data on wildtype cells.*

```
    union, unique, unsplit, which.max, which.min

Loading required package: S4Vectors

Attaching package: 'S4Vectors'

The following objects are masked from 'package:dplyr':

    first, rename

The following object is masked from 'package:tidyr':

    expand

The following objects are masked from 'package:base':

    expand.grid, I, unname

Loading required package: IRanges

Attaching package: 'IRanges'

The following objects are masked from 'package:dplyr':

    collapse, desc, slice

The following object is masked from 'package:purrr':

    reduce

Loading required package: GenomeInfoDb

Loading required package: AnnotationDbi

Loading required package: Biobase

Welcome to Bioconductor

    Vignettes contain introductory material; view with
    'browseVignettes()'. To cite Bioconductor, see
    'citation("Biobase")', and for packages 'citation("pkgname")'.
```

```
Attaching package: 'Biobase'


The following object is masked from 'package:MatrixGenerics':

    rowMedians


The following objects are masked from 'package:matrixStats':

    anyMissing, rowMedians



Attaching package: 'AnnotationDbi'


The following object is masked from 'package:dplyr':

    select


Loading required package: Biostrings

Loading required package: XVector


Attaching package: 'XVector'


The following object is masked from 'package:purrr':

    compact



Attaching package: 'Biostrings'


The following object is masked from 'package:base':

    strsplit


Loading required package: rtracklayer

Type 'citation("pROC")' for a citation.


Attaching package: 'pROC'


The following objects are masked from 'package:IRanges':
```

```
    cov, var


The following objects are masked from 'package:S4Vectors':

    cov, var


The following object is masked from 'package:BiocGenerics':

    var


The following objects are masked from 'package:stats':

    cov, smooth, var
Warning: replacing previous import 'GenomicRanges::union' by 'dplyr::union' when
loading 'Sierra'
Warning: replacing previous import 'GenomicRanges::intersect' by
'dplyr::intersect' when loading 'Sierra'
Warning: replacing previous import 'GenomicRanges::setdiff' by 'dplyr::setdiff'
when loading 'Sierra'
```

Set color palette.

```
res_colors <-
  c(IDR = "tan",
    gIDR = "#30B7BC", # bright teal
    eCV = "#AF275F", # light magenta
    mIDR = "#DE653A" # medium teal
  )
```

Download genome.

```
wget https://ftp.ebi.ac.uk/pub/databases/gencode/Gencode_human/release_41/GRCh38.p13.genome.fa.gz
gunzip GRCh38.p13.genome.fa.gz
```

Download GTF file.

```
wget https://ftp.ebi.ac.uk/pub/databases/gencode/Gencode_human/release_41/gencode.v41.chr_patch_hapl_sca
gunzip gencode.v41.chr_patch_hapl_scaff.annotation.gtf.gz
```

Upload eCLIP data and keep only chromosome one for faster computations.

```
(eclip_data <-
  read_tsv(
    file = "RBFOX2_new_reagents.peak_table.tsv",
    col_types = cols()) %>%
   dplyr::filter(chrom == "chr1"))
New names:
* `` -> `...1`
# A tibble: 25,391 x 13
```

```
    ...1 chrom  start    end strand IP_num_QC012b_5_S5_R1_001
   <dbl> <chr>  <dbl>  <dbl> <chr>                      <dbl>
 1     0 chr1   16239  16316 -                              2
 2     1 chr1   17435  17527 -                              9
 3     2 chr1  185698 185733 -                              1
 4     3 chr1  186700 186754 -                              1
 5     4 chr1  186866 186925 -                              1
 6     5 chr1  187457 187537 -                              2
 7     6 chr1  188049 188114 -                              4
 8     7 chr1  189416 189524 -                              3
 9     8 chr1  190076 190138 -                              1
10     9 chr1  190810 190867 -                              2
# i 25,381 more rows
# i 7 more variables: IP_num_QC012b_6_S6_R1_001 <dbl>,
#   IP_num_QC012b_7_S7_R1_001 <dbl>, IP_num_QC012b_8_S8_R1_001 <dbl>,
#   input_num_QC012b_5_S5_R1_001 <dbl>, input_num_QC012b_6_S6_R1_001 <dbl>,
#   input_num_QC012b_7_S7_R1_001 <dbl>, input_num_QC012b_8_S8_R1_001 <dbl>
```

Extract peaks intensities.

```
eclip_inten <-
  eclip_data %>%
  dplyr::select(contains("num_QC"))
```

Get PCA of peak intensities.

```
eclip_inten <- eclip_inten %>% ceiling()
pca <- princomp(log2(eclip_inten + 1))
ggplot(pca$loadings[1:8,TRUE] %>%
         as.data.frame() %>%
         rownames_to_column("Sample"),
       aes(x=Comp.1, y=Comp.2)) +
  ggrepel::geom_label_repel(aes(label=Sample),size=2) + geom_point() +
  theme_bw() + ggtitle("PCA biplot of peaks intensities")
```

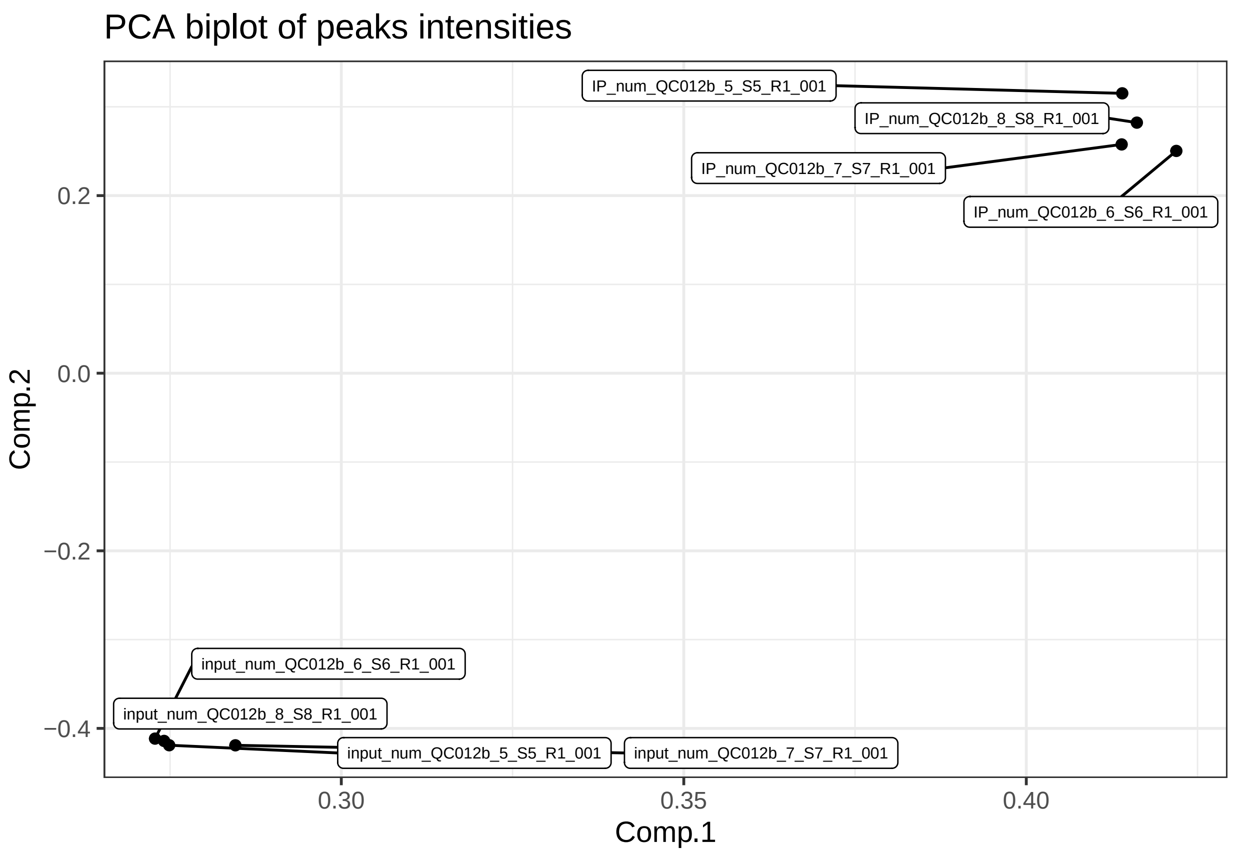


Get normalized IP intensities.

```
eclip_inten <- (eclip_inten %>% dplyr::select(contains("IP_num")) %>%
                 mutate_all(~ . + 1)) /
  (eclip_inten %>% dplyr::select(-contains("IP_num")) %>% mutate_all(~ . + 1))
```

Filter peaks with normalized intensity higher than 1.

```
tmp <- rowSums(eclip_inten >=
                 1) > 0
eclip_data <- eclip_data[tmp,]
eclip_inten <- eclip_inten[tmp,]
```

```
eclip_inten <-
      eclip_inten %>%
              mutate_all(~ . + 1.1) %>%
               as.matrix() %>%
  log()
```

Create subset with different number of replicates.

```
# Subset of j replicates.
eclip_inten_reps <-
  lapply(c(2, 3, 4), function(j) eclip_inten[,seq_len(j)])
```

```
names(eclip_inten_reps) <- paste0("n_rep=", c(2,3,4))

# Check dimensions.
lapply(eclip_inten_reps, dim)
$`n_rep=2`
[1] 24703     2

$`n_rep=3`
[1] 24703     3

$`n_rep=4`
[1] 24703     4
```

Set initial values for IDR, gIDR, and mIDR and number of iterations for eCV.

```
# Set parameters for each model.
methods_params <- list(
  eCV = list(max.ite = 1e4),
  IDR = list(
    mu = 2.5,
    sigma = 1,
    rho = 0.8,
    p = 0.5,
    eps = 1e-3,
    max.ite = 100
  ),
  gIDR = list(
    mu = 2.5,
    sigma = 1,
    rho = 0.8,
    p = 0.5,
    eps = 1e-3,
    max.ite = 100
  ),
  mIDR = list(
    mu = 2.5,
    sigma = 1,
    rho = 0.8,
    p = 0.5,
    eps = 1e-3,
    max.ite = 100
  )
)
```

### Data analysis.

#### FOX2 motif enrichment.

Create peak IDs.

```
eclip_data <-
  eclip_data %>%
  mutate(ID = paste0(chrom,":",start,"-",end,strand),
         score = 0)
```

Export table as a bed file.

```
eclip_data %>%
  dplyr::select(chrom, start, end, ID, score, strand) %>%
  write_tsv(file = "eclip_data.bed", col_names = FALSE)
```

Detect peaks matching FOX2 binding motif with Homer.

```
bin/findMotifsGenome.pl "eclip_data.bed" \
      GRCh38.p13.genome.fa \
      .aux \
      -find FOX2_all.motif -rna -size 75 \
      > "eclip_data_homer_res.bed"
rm -f -r .aux
```

Load Homer's results.

```
eclip_data <-
  eclip_data %>%
  mutate(motif=eclip_data$ID %in%
           read_delim(file = "eclip_data_homer_res.bed",
                      col_types = cols())$PositionID)
```

### Features distribution.

```
gtf <-
  import.gff("gencode.v41.chr_patch_hapl_scaff.annotation.gtf")
```

make TxDb.

```
txdb <-
  makeTxDbFromGFF("gencode.v41.chr_patch_hapl_scaff.annotation.gtf")
Import genomic features from the file as a GRanges object ... OK
Prepare the 'metadata' data frame ... OK
Make the TxDb object ...
Warning in .get_cds_IDX(mcols0$type, mcols0$phase): The "phase" metadata column contains non-NA values [illegible]
  stop_codon. This information was ignored.
OK
```

```
feature_dist <-
  eclip_data %>%
  makeGRangesFromDataFrame() %>%
  annotate_gr_from_gtf(gtf_gr = gtf,
                       gtf_TxDb = txdb,
```

```
                      transcriptDetails = TRUE)

Annotating 3' UTRs
Annotating 5' UTRs
Annotating introns
Annotating exons
Annotating CDS
```

#### Correlation analysis of impossed reproducible features.

```
tmp <- eclip_data$motif & (feature_dist$UTR3 == "YES" | feature_dist$intron == "YES")

R<- cor(eclip_inten[tmp, ])
r <- mean(R[upper.tri(R)])

CorCI(r,sum(tmp))
      cor    lwr.ci    upr.ci
0.7195490 0.6988817 0.7390157
```

#### Assess reproducibility with each method.

Create a table with all combinations of parameters.

```
(par_settings <- expand.grid(n_rep =  paste0("n_rep=",c(4,3,2)),
                            method=c("gIDR","mIDR","IDR","eCV"),
                            stringsAsFactors = FALSE) %>%
  mutate(par_com = seq_along(n_rep)))
     n_rep method par_com
1  n_rep=4   gIDR       1
2  n_rep=3   gIDR       2
3  n_rep=2   gIDR       3
4  n_rep=4   mIDR       4
5  n_rep=3   mIDR       5
6  n_rep=2   mIDR       6
7  n_rep=4    IDR       7
8  n_rep=3    IDR       8
9  n_rep=2    IDR       9
10 n_rep=4    eCV      10
11 n_rep=3    eCV      11
12 n_rep=2    eCV      12
```

```
# Set parallel scheduler.
future::plan(multisession, workers = 4)
perf_res <- NULL

for (i in par_settings$par_com) {
  set.seed(42)
  print(par_settings[i,])
  n_rep <- par_settings$n_rep[i]
  method <- par_settings$method[i]
```

```
  if(method != "eCV") {
    X <- preprocess(eclip_inten_reps[[n_rep]],
                         value_transformation = "identity",
                         jitter=1e-4)
  } else {
    X <- eclip_inten_reps[[n_rep]]
  }

  rep_index <- mrep_assessment(
          x = X,
          method = method,
          param = methods_params[[method]],
          n_threads = 4
        )$rep_index

   tmp <-  eclip_data$motif & (feature_dist$UTR3 == "YES" | feature_dist$intron == "YES")

   print(perf <- roc(tmp, rep_index, quiet = TRUE))
   perf_thr <- ci.coords(perf,conf.level=0.90,
                         x =  1/(1 + exp(-c(1:20) + 10)),
                         ret=c("threshold","tpr","tnr"))
    perf_thr <- rbind(perf_thr$tpr %>%
    as.data.frame() %>%
    mutate(threshold= 1/(1 + exp(-c(1:20) + 10)),
           perf="TPR (CI)") ,
        perf_thr$tnr %>%
    as.data.frame() %>%
    mutate(threshold= 1/(1 + exp(-c(1:20) + 10)),
           perf="TNR (CI)"))
  perf_thr$n_rep <- n_rep
  perf_thr$method <- method
  perf_res <- rbind(perf_res, perf_thr)
}

# Close me buddies.
future::plan(sequential)

# Save results.
saveRDS(perf_res,file="perf_resRealRBP.rds")
```

Arrange results for figure creation.

```
p <- perf_res %>%
mutate(n_rep = paste0("r=",str_remove(n_rep,"n_rep="))) %>%
  ggplot(aes(x=threshold,y=`50%`,color=method)) +
  facet_grid(perf~n_rep,switch ="y") +
   geom_ribbon(color = NA,
    aes(x = threshold, ymin = `5%`, ymax = `95%`,fill=method),
     alpha = 0.3)  +
   geom_line() +
  scale_color_manual(values=alpha(res_colors, 1)) +
  scale_linetype_manual(values=c("miR1"="solid", "miR124"="dashed")) +
  scale_fill_manual(values=alpha(res_colors, 0.6)) +
```

```
    scale_y_continuous(position = "right")+
   theme_bw() + theme(
                    legend.title = element_text(size=9),
                    legend.text = element_text(size=7, face = "italic"),
                    strip.background = element_rect(fill=alpha("gray", 0.25)),
                 legend.background = element_rect(fill = alpha("white", 0.1))) +
   guides(color=guide_legend(ncol=1,override.aes = list(size = 2))) +
   labs(y="",fill="Method",linetype="Transfection",
       color="Method",x="Reproducibility threshold (%)") +
   scale_x_continuous(breaks=c(0, 0.25, 0.5, 0.75, 1),
                    labels = c("0","25","50","75","100"))
p
```

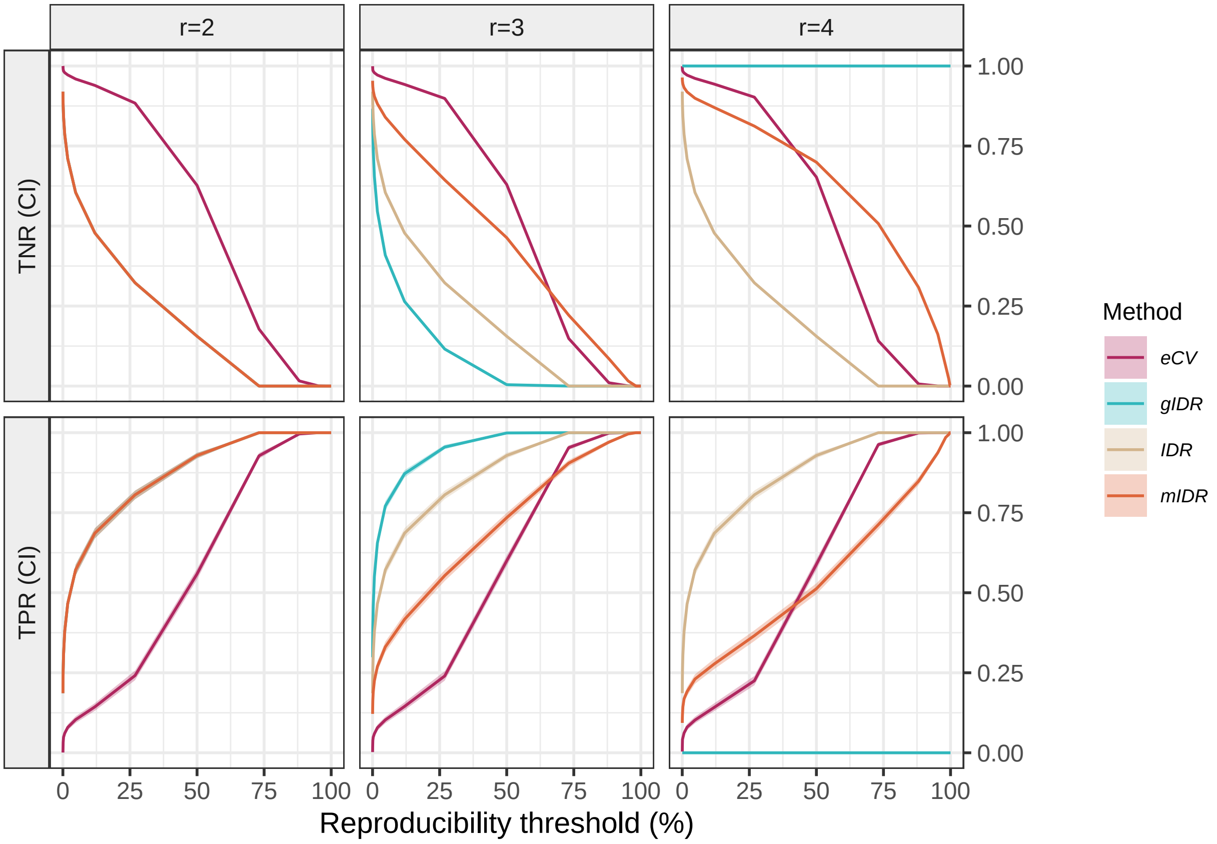


```
ggsave(filename = "Figure6.tiff",plot = p,device = "tiff",
       dpi=300,units = "in",width = 7,height = 5,scale = 0.85)
```
